# TCDD dysregulation of lncRNA expression, liver zonation and intercellular communication across the liver lobule

**DOI:** 10.1101/2023.01.07.523119

**Authors:** Kritika Karri, David J. Waxman

## Abstract

The persistent environmental aryl hydrocarbon receptor agonist and hepatotoxin TCDD (2,3,7,8-tetrachlorodibenzo-*p*-dioxin) induces hepatic lipid accumulation (steatosis), inflammation (steatohepatitis) and fibrosis. Thousands of liver-expressed, nuclear-localized lncRNAs with regulatory potential have been identified; however, their roles in TCDD-induced hepatoxicity and liver disease are unknown. We analyzed single nucleus (sn)RNA-seq data from control and chronic TCDD-exposed mouse liver to determine liver cell-type specificity, zonation and differential expression profiles for thousands of IncRNAs. TCDD dysregulated >4,000 of these lncRNAs in one or more liver cell types, including 684 lncRNAs specifically dysregulated in liver non-parenchymal cells. Trajectory inference analysis revealed major disruption by TCDD of hepatocyte zonation, affecting >800 genes, including 121 IncRNAs, with strong enrichment for lipid metabolism genes. TCDD also dysregulated expression of >200 transcription factors, including 19 Nuclear Receptors, most notably in hepatocytes and Kupffer cells. TCDD-induced changes in cell–cell communication patterns included marked decreases in EGF signaling from hepatocytes to non-parenchymal cells and increases in extracellular matrix-receptor interactions central to liver fibrosis. Gene regulatory networks constructed from the snRNA-seq data identified TCDD-exposed liver network-essential lncRNA regulators linked to functions such as fatty acid metabolic process, peroxisome and xenobiotic metabolic. Networks were validated by the striking enrichments that predicted regulatory IncRNAs showed for specific biological pathways. These findings highlight the power of snRNA-seq to discover functional roles for many xenobiotic-responsive lncRNAs in both hepatocytes and liver non-parenchymal cells and to elucidate novel aspects of foreign chemical-induced hepatotoxicity and liver disease, including dysregulation of intercellular communication within the liver lobule.

## Introduction

2,3,7,8-Tetrachlorodibenzo-*p*-dioxin (TCDD) is a persistent hepatotoxic environmental chemical that directly binds to aryl hydrocarbon receptor (AhR), a ligand-activated transcription factor that dysregulates hundreds of genes ^1, 2^. TCDD-activated AhR translocates to the nucleus, where it heterodimerizes with Arnt and then binds to chromatin at thousands of regulatory sequences controlling transcription of linked target genes ^3^. TCDD also induces AhR-dependent nongenomic signaling via changes in intracellular Ca^2+^ and activation of the tyrosine kinase src and downstream kinases ^4, 5^. TCDD exposure leads to a broad range of hepatotoxic and other effects in both mice and humans ^6^, including immune suppression, hepatic lipid accumulation and progression to non-alcoholic steatohepatitis (NASH) with fibrosis ^7-9^. TCDD also has the potential to induce epigenetic alterations that may be passed along trans-generationally ^10^. Many metabolic and other pathways dysregulated by TCDD have been identified ^11^, but the precise mechanisms of TCDD action and in particular the role of AhR downstream genes that mediate the effects of chronic TCDD exposure on liver metabolic diseases, including NASH and liver fibrosis, remains an area of active investigation.

The liver carries out a wide array of physiological processes, including energy homeostasis, gluconeogenesis, serum protein synthesis and xenobiotic metabolism. These diverse processes are enabled by the liver’s complex architecture, where hepatocytes and non-parenchymal cells (NPCs) are organized within local repeating structures known as the liver lobule. Hepatocytes constitute about 70% of the liver mass while the remaining 30% consists of NPCs, primarily endothelial cells, hepatic stellate cells, and various immune cell populations, including Kupffer cells (liver resident macrophages) ^12^. Single-cell transcriptomics has the power to elucidate cell-specific responses and spatial zonation ^13, 14^ and has transformed our understanding of the roles these diverse cells play in liver homeostasis ^15^ and pathophysiology ^16, 17^, including following chronic TCDD exposure ^18^.

Long non-coding RNAs (lncRNAs) are RNAs longer than 200 bp with little or no protein-coding potential. Thousands of liver-expressed IncRNAs are nuclear enriched in the nucleus where many are tightly bound to liver chromatin ^19^, which enables them to play essential regulatory roles in modulating chromatin function and gene expression ^20^. Several hundred liver lncRNAs show significant differences in expression between males and females ^19, 21, 22^ and/or respond to drug and foreign chemical exposure ^19, 23-25^. Individual lncRNAs have been linked to hepatotoxicity induced by foreign chemical exposures ^26, 27^, however, a global investigation of the roles of lncRNAs in foreign chemical-induced hepatotoxicity, and in particular TCDD-induced hepatoxicity, is lacking. Recently, Nault *et al* used snRNA-seq to characterize liver cell type-specific transcriptional changes in a chronic (28 day) TCDD exposure model and reported widespread dysregulation of gene expression in multiple liver cell types, as well as marked expansion of the liver macrophage population and extensive TCDD-induced changes in hepatic zonation ^18, 28^. However, the effects of TCDD on the thousands of liver-expressed lncRNAs recently identified and shown to be amenable to analysis using snRNA-seq ^22^ were not considered.

Here, we utilized a single cell transcriptome reference gene set comprised of 76,011 mouse genes, including 48,261 mouse liver-expressed lncRNAs whose gene structures and isoform patterns we recently characterized ^22^, to discover lncRNAs whose expression and zonation across the liver lobule is dysregulated in either hepatocytes or liver NPCs following chronic TCDD exposure. We also characterize the liver cell type-dependent effects of TCDD on expression of more than 1,500 transcription factors, including AhR itself and many members of the nuclear receptor (NR) superfamily. Furthermore, we map the global landscape of intercellular signaling between liver cell types and explore TCDD-induced changes in intrahepatic cell-cell communication, which plays a critical role in tissue homeostasis and injury response ^29^ and may contribute to TCDD hepatotoxicity. Finally, we construct gene regulatory networks to link individual TCDD-responsive lncRNAs to biological functions, and using network centrality metrics, we identify network-essential regulatory nodes ^30, 31^ that provide systems-level insight into the regulatory functions of both lncRNAs and protein-coding genes contributing to the widespread hepatic effects of TCDD exposure.

## Methods

### Data processing of snRNA-seq samples

Single nucleus-based RNA-seq raw sequencing data (Fastq files) for control and TCDD-exposed male mouse livers ^18^ were downloaded from GEO (accession #GSE148339; https://www.ncbi.nlm.nih.gov/geo/), processed using CellRanger (v3.1.0) ^32^ and aligned to the mouse mm10 reference genome. Mapped snRNA-seq reads were assigned to individual genes using a custom GTF file (see below) whose entries span the full gene body of each gene (rather than exonic regions only, as is typically carried out when counting single cell-RNA-seq reads) to enable counting of intronic reads, which comprise a substantial fraction of the snRNA-RNA sequencing reads. For gene pairs that overlap each other on the same strand (whose gene counts would be excluded by the 10X Genomics CellRanger’s counting algorithm across the entire region of overlap), we modified the GTF file entries to include all exons of both overlapping genes, plus all non-exonic gene regions that were unique to each gene, as detailed elsewhere ^22^. Thus, sequence reads that map to an exon of one gene but overlap an intronic region of the other, overlapping gene were uniquely counted for the first gene by excluding the overlapping intronic region of the second gene from the GTF annotation file for that gene.

The custom GTF file used for counting (GTFB_FullGeneBody_MouseLiver_snRNAseq.gtf) is available at https://tinyurl.com/GTF-MouseLiver48kLncRNAs and includes a total of 76,011 mouse mm10 genes, plus 91 ERCC spike-in control sequences, as follows: 20,973 RefSeq protein-coding genes including 13 mitochondrial genes; 2,077 RefSeq non-coding genes (i.e., RefSeq genes assigned NR accession numbers that do not overlap the set of 48,261 lncRNAs described immediately below); and a total of a total of 52,961 lncRNA genes (48,261 liver-expressed lncRNAs ^22^ and 4,700 other lncRNA genes, namely, 4,697 Ensembl noncoding lncRNAs that do not overlap the RefSeq NR dataset or the set of 48,261 liver-expressed lncRNAs, and 3 other lncRNAs (lnc-LFAR1, LeXis, Lnclgr) that were absent from the above lists). (Of note, the set of 4,697 Ensembl genes included 3,176 genes designated ‘lincRNA’ and 1,521 genes designated ‘anti-sense’ with respect to a protein-coding gene in the Supplemental tables). Human and rat lncRNA sequences orthologous to the above sets of mouse lncRNAs were identified using methods described previously ^24^ and are included in Table S1B.

### Integration and clustering

Feature-barcode matrices were generated using the 10x Genomics Cell Ranger pipeline (v3.1.0) ^32^. snRNA-seq data from the control and TCDD-exposed liver samples were combined using the CellRanger *aggr* command. The resulting aggregated matrix was processed using Seurat package (v3.0) ^33^ in R (v.3.6.0). Cells with fewer than 200 genes detected, less than 400 UMI per cell, or >5% mitochondrial contamination were excluded. Doublets and multiplets were identified and removed from the single-cell sequencing data using scDblFinder ^34^, a doublet detection algorithm that uses several methods, including artificial doublet generation, parameter optimization and thresholding (with default parameters). The final count matrix used for clustering included 15,573 nuclei (9,597 nuclei from control liver; 5,976 nuclei from TCDD-exposed liver).

The integrated count matrix data from Cellranger was normalized by dividing the UMI count per gene by the total UMI count in the corresponding cell and then log-transforming the data. Highly variable genes were identified using the *FindVariableGenes* function of Seurat (v3) with default parameters. Variable genes were projected onto a low-dimensional space using principal component (PC) analysis, with the number of PCs set to 10 based on an inspection of elbow plots of the variance explained for each dataset. Batch correction was performed using Harmony ^35^ using the Seurat object as input, with default parameters to remove from the embedding the influence of dataset-of-origin factors. We input to Harmony normalized gene matrix files saved as a Seurat object along with pre-calculated PC analysis embedding based on 10 PCs using default parameters. Graph-based clustering of the cells based on their gene expression profiles was implemented using the *FindClusters* function in Seurat. The clusters obtained were visualized in Seurat v3 using the UMAP function (resolution: 0.2 and PC=10). Cell identities were assigned to each cluster based on the expression of established marker genes (Fig. S1B).

### Differential expression analysis

Differential gene expression between cell clusters under a given set of biological conditions (e.g., periportal vs pericentral hepatocytes from control liver), or between biological conditions for a given cell cluster (e.g., control endothelial cells vs TCDD-exposed endothelial cells) was performed using the Locally Distinguishing function of the 10x Genomics Loupe Browser (v.6.0) (see Loupe Browser file in Supplement). This method implements the negative binomial test based on the sSeq method ^36^ with Benjamini-Hochberg correction for multiple tests; it uses log-normalized average expression values and the distribution of UMIs across the two specific cell populations being compared to obtain fold-change and FDR values for each pair-wise gene expression comparison. Genes were identified as showing significant differential expression between two cell populations if they met the stringent threshold of |fold-change| >4 at FDR <0.05.

### Functional enrichment analysis

Enrichment analysis of protein-coding gene lists from various outputs was performed using DAVID ^37^ with default parameters, except that GO FAT terms were used in place of GO DIRECT terms to include a broader range of enrichment terms than the default GO DIRECT option. We used a custom script prepared by Dr. Alan Downey-Wall of this laboratory (https://github.com/adowneywall/Davidaggregator) to aggregate and reformat DAVID output files from multiple gene lists, with each row presenting top enriched annotation clusters and individual columns presenting enrichment score, p-value, FDR and other such data. Top terms from each DAVID annotation cluster having a cluster enrichment score > 3 and top FDR < 0.05 were used in downstream analysis.

### TCDD-elicited perturbations in hepatocyte zonation

Monocle2 ^38^ was used to infer spatial trajectories for hepatocyte nuclei from control and TCDD-exposed liver. Genes showing significant zonation in control hepatocytes at q-value <0.001 were identified using generalized linear models via the *differentialGeneTest* function in Monocle. Genes that were differentially zonated between control liver and TCDD-exposed liver were identified by deriving a common pseudotime trajectory using Monocle2. We used the *conditionTest* function implemented in the package tradeSeq in Bioconductor ^39^ to compare the two conditions (control and TCDD exposure) along a common trajectory to detect gene expression changes indicative of differential progression. The *conditionTest* function tests the null hypothesis that genes have identical expression patterns in each condition across a *pseudotime*. We extracted perturbed genes that were significant at FDR < 0.001 and created matched heatmaps for both control and TCDD-exposed liver using the *pheatmap* function of Monocle2. Heatmaps were further clustered individually for each condition to assign the zonation labels shown in Table S2C.

### NR and TF expression analysis

The Dotplot function in Seurat (v3) was used to present expression profiles for each control mouse liver cell cluster for all 49 mouse nuclear receptor (NR) genes and for AhR. Differential expression analysis between control and TCDD-exposed liver was carried out for those 50 genes, and also for a much larger listing of 1,533 mouse TFs (Table S3B) downloaded from AnimalTFDB3.0 (http://bioinfo.life.hust.edu.cn/AnimalTFDB/) ^40^ by using the Locally Distinguishing function of the 10x Genomics Loupe Browser (v.6.0) (Table S3C), as described above. Fisher’s exact test was used to identify those TF families whose gene members were significantly enriched (at p-value <0.05) in the set of differentially expressed TFs from TCDD-exposed liver (Table S3D).

### Cell-cell communicaton analysis

CellChat ^29^ inputs gene expression data, assigns cell labels as input, and then models cell-cell communication probabilities by integrating gene expression with prior knowledge of the interactions between signaling ligands, receptors and their cofactors. We input to CellChat processed transcriptomic data from the 15,573 nuclei from control liver, and from TCDD-exposed liver, together with cell labels extracted from the Seurat object. Next, we inferred cell-cell communication probabilities using the *computeCommunProb* function, which infers biologically significant interactions by assigning each interaction a probability value and then performs a permutation test. We ran CellChat on the control liver and the TCDD-exposed liver datasets separately and then merged the CellChat objects from each analysis using the *mergeCellChat* function. This resulted in a total of 1,087 interactions (225 unique interactions) in control and TCDD-exposed liver (Table S4A). These interactions were grouped into 52 signaling pathways, as defined by CellChatDB (Fig. 5B). Pathways that were either conserved between biological conditions or were specific to TCDD-exposed liver were identified by comparing the information flow for each signaling pathway, which is defined by the sum of communication probabilities among all pairs of cell groups in the inferred network (i.e., the total weights in the network). Cell-cell interaction networks were visualized using the circle plot layout in CellChat, with edge colors consistent with the cell sources, and with edge weights proportional to the interaction strength (i.e., using a thicker edge to indicate a stronger signal).

### Gene regulatory network analysis

Gene regulatory networks for control liver and for TCDD-exposed liver were inferred by analysis of the snRNA-seq data using bigSCale2 ^30^. Nuclei from control liver and from TCDD-exposed liver were split into two Seurat objects and passed to bigSCale2 ^30^, where each network was constructed using the normal (default) clustering parameter, with granularity set to the highest setting and applying an edge cutoff an edge cutoff corresponding to the top 99.9 quantile for correlation coefficient. bigSCale2 retains edges that are expected to represent actual regulatory links by removing genes that do not have a direct edge with a known GO regulator (GO sub-setting step). This was achieved using a list of GO regulators consisting of the union of the genes in GO:0006355 “regulation of transcription, DNA-templated” (N=2,776) and GO:000370 “DNA-binding transcription factor activity” (N=1,001). We also included our list of 52,961 lncRNAs (see above) as potential regulators in the GO sub-setting step. Network specifications are shown in Table S5G. The networks obtained were converted into json files for visualization by Cytoscape ^41^ using the *toCytoscape* function of the package iGraph (v1.3.5) (https://igraph.org/r) in R. Specifically, the network json file was imported in Cytoscape using “*Import Network from File”*. Next, we used “*forced-directed”* layout in Cytoscape, where each network’s layout was derived from 10,000 iterations of the Fruchterman-Reingold algorithm ^42^ and using the Nogrid parameter (seed=7). The resultant networks and derived subnetworks shown in the various figures are available are available at Ndexbio.org at https://tinyurl.com/snTCDDliverNetworksKarriWaxman. Gene modules in each network were discovered using the Glay community cluster algorithm ^43^ in Cytoscape ^41^, whereby the overall network topology was used to subdivide the overall networks generated by bigSCale2 into functional modules.

Genes in each module were input to DAVID ^37^ for gene ontology enrichment analysis. Node rank was calculated for each of four key bigSCale2 network metrics (Betweenness, Closeness, Degree, PAGERANK). Protein-coding genes that ranked within the top 100 nodes for any one of the above four network metrics were deemed to be network-essential nodes that identify regulatory protein-coding genes; and lncRNAs that ranked within the top 50 nodes for any one of the above four network metrics were deemed to be network-essential nodes corresponding to regulatory lncRNAs (Table S5B)

### Master regulators

To identify master regulators, we extracted a subnetwork compromised of the network-essential genes, i.e., the top 100 ranked nodes for protein-coding genes, and the top 50 ranked nodes for lncRNAs, evaluated separately for each of the 4 network metrics, and then recalculated network metrics. The critical nodes in these subnetworks were identified using five network metrics: Stress, Degree, Betweenness, Closeness and Neighborhood Connectivity, and were calculated using the “*Analyze Network*” option in Cytoscape. We then designated as master regulators, i.e., critical nodes regulating the regulators, the top 5 ranked nodes, considering each of the 5 metrics in the ranking independently (Table S5B).

### IPA Analysis

We used Ingenuity Pathway Analysis software (http://www.ingenuity.com) to validate the regulatory role of protein-coding gene master regulators from the control liver and from the TCDD-exposed liver networks. Putative gene targets of protein-coding gene master regulators were provided as input to IPA. IPA core analysis of these gene lists yielded information on the upstream regulators, top enriched canonical pathways, diseases, and toxicological functions. The resulting data output by IPA for all protein-coding gene master regulators is included in Table S5E.

### Enrichment heatmap

Protein-coding genes that made direct connections to a regulatory lncRNAs in a bigSCale2 network were input to Metascape ^44^. We performed multi-list comparative analysis within and across the networks to identify pathways that are shared between or are specific to a lncRNA’s gene targets. The clustered heatmaps obtained depict -log10 P-value of top enriched pathways across multiple lncRNAs target lists.

## Results

### Cell-type specific responses of liver lncRNAs to TCDD exposure

We integrated and clustered snRNA-seq datasets representing a total of 15,573 nuclei extracted from livers of mice exposed to TCDD chronically (28 days), or vehicle control ^18^ (Table S1A). Eight major cell clusters representing 8 distinct liver cell subpopulations were identified using established liver cell type-specific marker genes (Fig. S1). Major treatment-dependent changes in the proportions of key liver cell types were found, including large decreases in the proportion of periportal hepatocytes and hepatic stellate cells, and increases in Kupffer cells, endothelial cells, B cells and T cells (Table S1A), consistent with ^18^. The snRNA-seq data was of high quality and sufficient sequencing depth for us to detect 33,756 lncRNA genes using a custom GTF file encompassing a total of 76,011 genes, including 52,961 lncRNA genes (see Methods).

Differential expression analysis applied to each liver cell cluster identified a total of 7,821 genes, including 4,016 lncRNA genes, that were dysregulated following TCDD exposure in at least one liver cell type at an expression |fold-change|> 4 and FDR <0.05 (Table S1B). The largest number of TCDD gene responses was seen in pericentral hepatocytes, where AhR is most highly expressed, followed by periportal hepatocytes and then Kupffer cells (Fig. 1, Table S1A), where AhR levels are much lower (see below). In hepatocytes, many more genes were induced by TCDD than were repressed, both for protein-coding genes and for lncRNAs, whereas in the NPC clusters gene down regulation was generally more extensive than gene up regulation (Fig. 1, Fig. S2). We observed large differences in the cell type-specificity of TCDD-induced gene responses for lncRNAs as compared to protein-coding genes. Thus, 75% of TCDD-responsive lncRNAs, but only 30% of TCDD-responsive protein-coding genes, showed a response to TCDD in hepatocytes but not in NPCs; whereas 54% of TCDD-responsive protein-coding genes, but only 17% of TCDD-responsive lncRNAs, responded to TCDD in one or more NPCs, but not in hepatocytes (Fig. S2D).

**Fig. 1.**
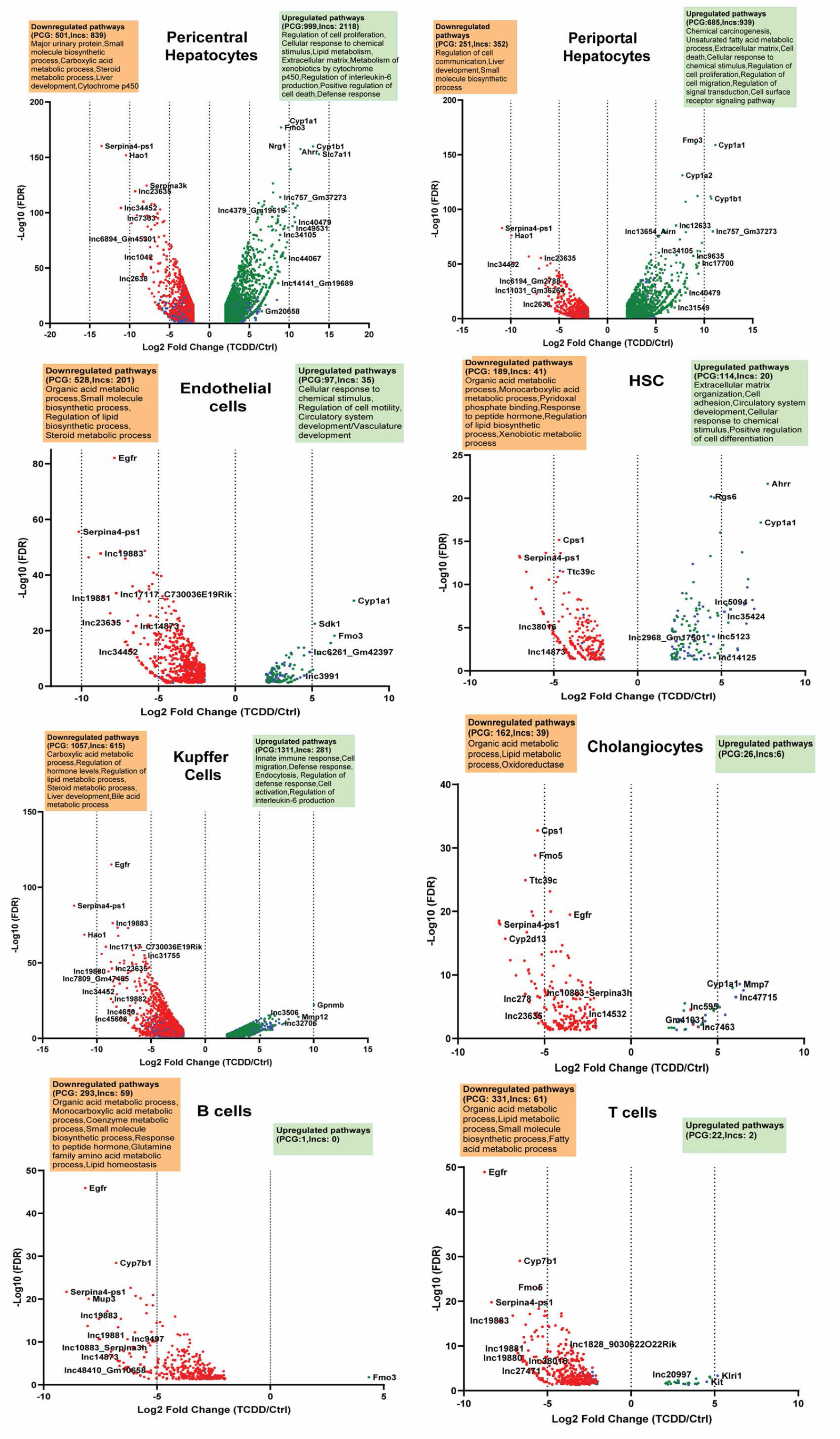
TCDD-elicited gene responses in individual liver cell populations. Volcano plots showing protein-coding genes and lncRNA genes whose expression is dysregulated by TCDD at |fold-change| >4 and FDR <0.05 threshold in each of the 8 indicated liver cell clusters. See Table S1B for full gene listing.

Top TCDD-responding protein-coding genes include classic AhR target genes, such as Cyp1a1 and Cyp1a2 (Fig. S3A). In hepatocytes, the most highly inducible lncRNAs included lnc17700* and Gm2968 (lnc19317) (Fig. S3B), with fold-change values > 400 (Table S1B). Well-characterized lncRNAs induced by TCDD in both pericentral and periportal hepatocytes include Snhg15 (lnc9442*), whose increased expression has been linked to liver metastasis and poor overall survival in colorectal cancer ^45^, and Lrrc75-as1 (lnc9790*), which inhibits cell proliferation and migration in colorectal cancer ^46^. In contrast, Lhx1os (lnc9944), which is anti-sense to the transcription factor Lhx1, was induced 100-fold by TCDD specifically in pericentral hepatocytes, while 4933401D09Rik (lnc13688) was induced 45-fold specifically in periportal hepatocytes (Table S1B). TCDD responses that were highly specific for individual NPC populations include the >30-fold up regulation of two adjacent novel lncRNAs (lnc47715, lnc47717) in TCDD-exposed cholangiocytes (Fig. S3A), and the >30-fold up regulation of a novel lncRNA (lnc5094) in hepatic stellate cells. Neat1 (lnc14746*), a positive regulator of liver fibrosis ^47^, was down regulated 18-fold in TCDD-exposed Kupffer cells. Thirteen lncRNAs, including lnc10883 and lnc17117, were significantly repressed by TCDD across all liver cell types (Table S1C, Fig. S3C).

Functional enrichment analysis of the set of TCDD-responsive protein-coding genes identified overlapping groups of pathways that were down regulated in multiple NPC cell types. For example, genes associated with lipid homeostasis and related terms were down regulated in endothelial cells, hepatic stellate cells, cholangiocytes and immune cells (B cells, T cells, Kupffer cells). Pathways related to xenobiotic metabolism, cell proliferation and cytokine production were strongly up regulated in hepatocytes. In contrast, extracellular matrix was most strongly induced in hepatic stellate cells, while immune response pathways showed very strong increases in Kupffer cells (Fig. 1, Table S1D).

### Zonated expression of lncRNAs in hepatocytes

Hepatocyte nuclei from healthy (control) mouse liver were resolved using the spatial inference algorithm Monocle2 ^38^ to identify 204 lncRNAs and 1,744 protein-coding genes (Table S2A) that showed significant zone-dependent expression in hepatocytes across the liver lobule (Table S2A). The functional enrichments obtained were consistent with prior findings using experimentally validated hepatocyte zonation markers ^13^, such as Gulo (pericentral) and Cyp2f2 (periportal) (Fig. 2A, Table S2B). Examples of strong pericentral hepatocyte zonation bias include lnc32377 (Cyp2c53-ps) and lnc34787 (Snhg11), which functions as a tumor promoter in hepatocellular carcinoma ^48^. Strong pericentral zonation was found for lnc2311 (Platr4), which ameliorates steatohepatitis in mouse liver ^49^, and for lnc14025 (Dreh), which promotes cell proliferation in hepatitis B virus-associated hepatocellular carcinoma ^50^. Comparing zonal distributions between control and TCDD-exposed hepatocyte nuclei revealed major disruption of hepatocyte zonation following TCDD exposure, with 810 genes, including 121 lncRNAs, showing differential expression along the spatial trajectory between the two conditions (Table S2C). Fig. 2B shows examples of disrupted zonation of protein-coding genes (Car8, Etnk1) and lncRNAs (Cyp2c53-ps (lnc32377), lnc39934) in TCDD-exposed hepatocytes. Top enrichments for the zone-dysregulated protein-coding genes included carboxylic acid metabolic process, lipid metabolic process, apoptotic process and peroxisome (Table S2D).

**Fig. 2.**
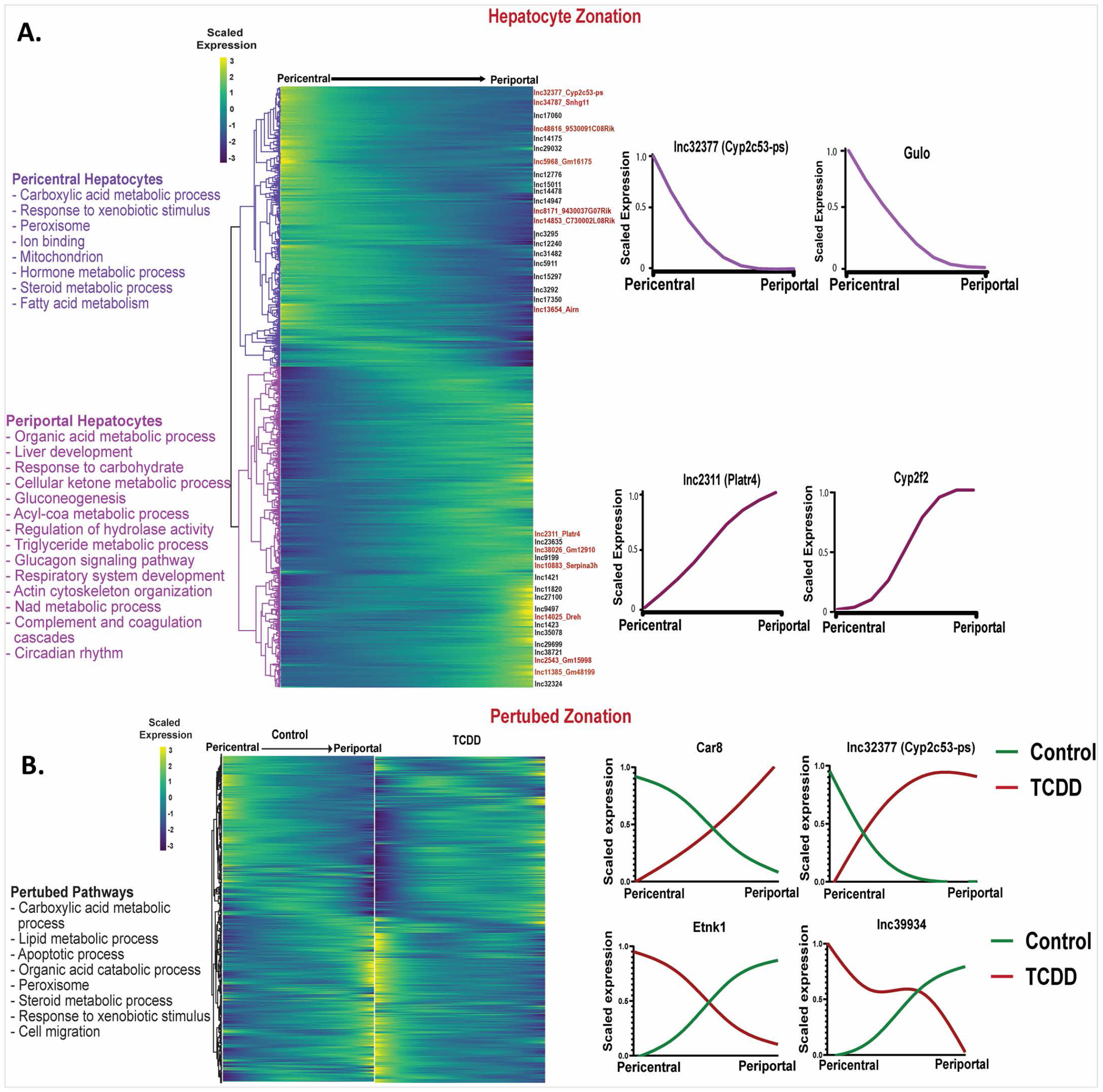
Dysregulation of hepatocyte zonation across the liver lobule in TCDD-exposed liver. **A**. Heatmap showing relative expression of 1,744 genes that show zonated expression in hepatocytes from healthy (control) mouse liver. Select zonated lncRNAs are marked at the right. Zonation profiles for select genes are shown at the far right and top enriched terms for each hepatocyte zone are listed at the left. **B**. Matched heatmaps for 810 genes that are differentially zonated between control and TCDD-exposed hepatocytes at FDR <0.001. See Table S2 for full datasets.

### Impact of TCDD on expression profiles for Nuclear Receptors (NR) and AhR

TFs from the NR superfamily are activated by steroids, fatty acids and other natural ligands, and in certain cases by foreign chemicals that dysregulate key liver functions, such as glucose and cholesterol metabolism, bile homeostasis and xenobiotic metabolism ^51, 52^. To better understand the potential for TCDD to alter these NR-dependent physiological processes and pathophysiological responses, we examined the expression profiles across liver cell types of all 49 mouse NRs ^53^, as well as that of Ahr, under both basal conditions and following TCDD exposure. The 49 NRs were grouped into five separate clusters based on their physiological functions ^54^, as marked in Fig. 3A. NRs in the bile acid and xenobiotic metabolism cluster and in the lipid metabolism and energy homeostasis cluster generally showed the highest expression (Fig. 3A). Expression levels were similar between periportal and pericentral hepatocytes, with the exception of three receptors with a strong pericentral bias: Nri13 (CAR), an important regulator of hepatic drug and xenobiotic metabolism and energy metabolism ^55^; Esrrg, a key regulator of hepatic gluconeogenesis ^56^; and AhR itself. Ppara, which along with Ppard and Pparg plays a critical role as a lipid sensor and regulator of lipid metabolism ^57, 58^, showed higher expression in both hepatocyte clusters than in the NPC clusters. Nr1h4 (FXRα), a bile acid sensor that regulates hepatic bile acid metabolism ^59^, showed a liver cell type specificity similar to Ppara, except that its expression was high in hepatic stellate cells, which have a major role in the deposition of extracellular matrix during liver fibrosis. Of note, Nr1h4 (FXRα) suppresses liver fibrosis development, and its deficiency is associated with fatty liver and increased susceptibility to NASH in high fat diet-fed mice ^60^. In contrast, expression of Nr1h5 (FXRβ), which is activated by lanosterol but not bile acids ^61^, was largely restricted to hepatic stellate cells. Rarb, which also showed high specificity for hepatic stellate cell expression, inhibits hepatic stellate cell activation in NAFLD ^62^. Finally, Rora, which decreases oxidative stress and has the ability to attenuate NAFLD progression ^63, 64^, was expressed in most liver cell types, with highest expression seen in cholangiocytes.

**Fig. 3.**
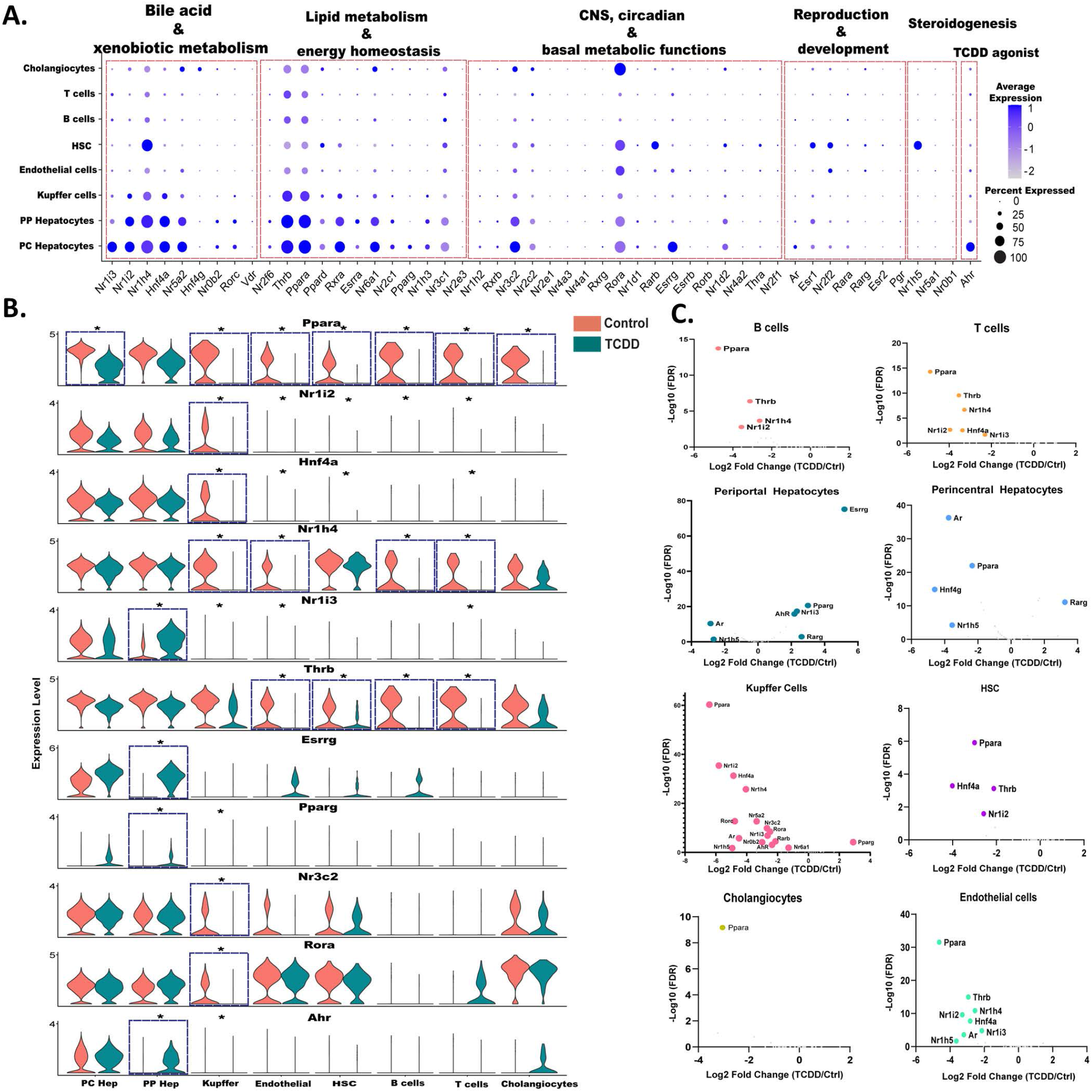
snRNA-seq expression profiles for AhR and members of the NR superfamily. **A**. Dot plot expression profiles for all 49 mouse NR genes, and for Ahr, in each of 8 mouse liver cell types from control mouse liver. **B**. Violin plots showing changes in expression of select NRs, and AhR, between control and TCDD-exposed mouse liver in the 8 liver cell types. Asterisk indicates significant difference in expression at FDR <0.05. **C**. Volcano plots depicting control versus TCDD liver differentially expressed NRs in each liver cell type. X-axis, log2 fold-change values for differential expression for control vs. TCDD within each cell cluster; Y-axis, - log10 (FDR) value to mark the significance of differential expression. Also see Table S3A.

TCDD altered the expression of AhR and of 19 out of the 49 NRs in at least one liver cell type. AhR and four NRs (Nr1i3 (CAR), Esrrg, Pparg, Rarg) were up regulated by TCDD in periportal hepatocytes. The increase in Nr1i3 (CAR) is consistent with a prior report in bulk liver analysis ^65^ and leads to a reversal of Nr1i3 (CAR) zonation (Table S2C). Similarly, the periportal increases in Esrrg and AhR largely abolish the pericentral zonation of those receptors seen in control hepatocytes (Fig. 3B, Table S2A). In addition, TCDD induced large decreases in expression of Ppara in all major liver cell types except periportal hepatocytes, which may contribute to TCDD disruption of PPARA-regulated lipid metabolism ^66^. In Kupffer cells, TCDD exposure led to widespread decreases in many NRs, including Ppara, Nr1i2 (PXR), Nr1h4 (FXRα) and Hnf4a. The down regulation of Hnf4a may contribute to TCDD-induced liver injury, insofar as Hnf4a repression occurs in several mouse models of acute liver damage, including sepsis and liver injury following high dose exposure to acetaminophen or carbon tetrachloride ^67, 68^. Thrb was down regulated by TCDD in both endothelial cells and hepatic stellate cells, as well as in B cells and T cells (Fig. 3B, Fig. 3C). Hepatic thyroid hormone signaling is crucial in the development and progression of NASH, and reduced expression of Thrb has been linked to NASH in human liver ^69^.

### Global evaluation of TF dysregulation in TCDD-exposed liver

Given the major changes in NR expression seen following TCDD exposure, above, we investigated more broadly the effects of TCDD on the expression of 1,533 TFs grouped into 72 TF families (Table S3B). TCDD induced or repressed the expression of 239 of these TFs (including 19 NRs discussed above) by > 4-fold in at least one liver cell type (Table S3C). Overall, 5 of the 72 families showing significant enrichment (p <0.05) for TFs being dysregulated by TCDD (Fig. S4). TF dysregulation was most widespread in hepatocytes and Kupffer cells (Fig. 4A). Nine TFs, including the NRs Ppara and Nr1i2, were dysregulated by TCDD in five or more cell types, all but one of which (Ahrr) were down regulated in multiple cell types (Fig. 4B). Examples include: Onecut1, a liver TF implicated in regulation of liver sex differences ^70^, consistent with the loss of liver sex differences following TCDD exposure ^71^; Mafb, which promotes macrophage polarization into an anti-inflammatory M2 state ^72^; and the lipogenic TF Mlxipl (ChREBP) ^73^, which was strongly down regulated in all six NPC populations, but not in hepatocytes.

**Fig. 4.**
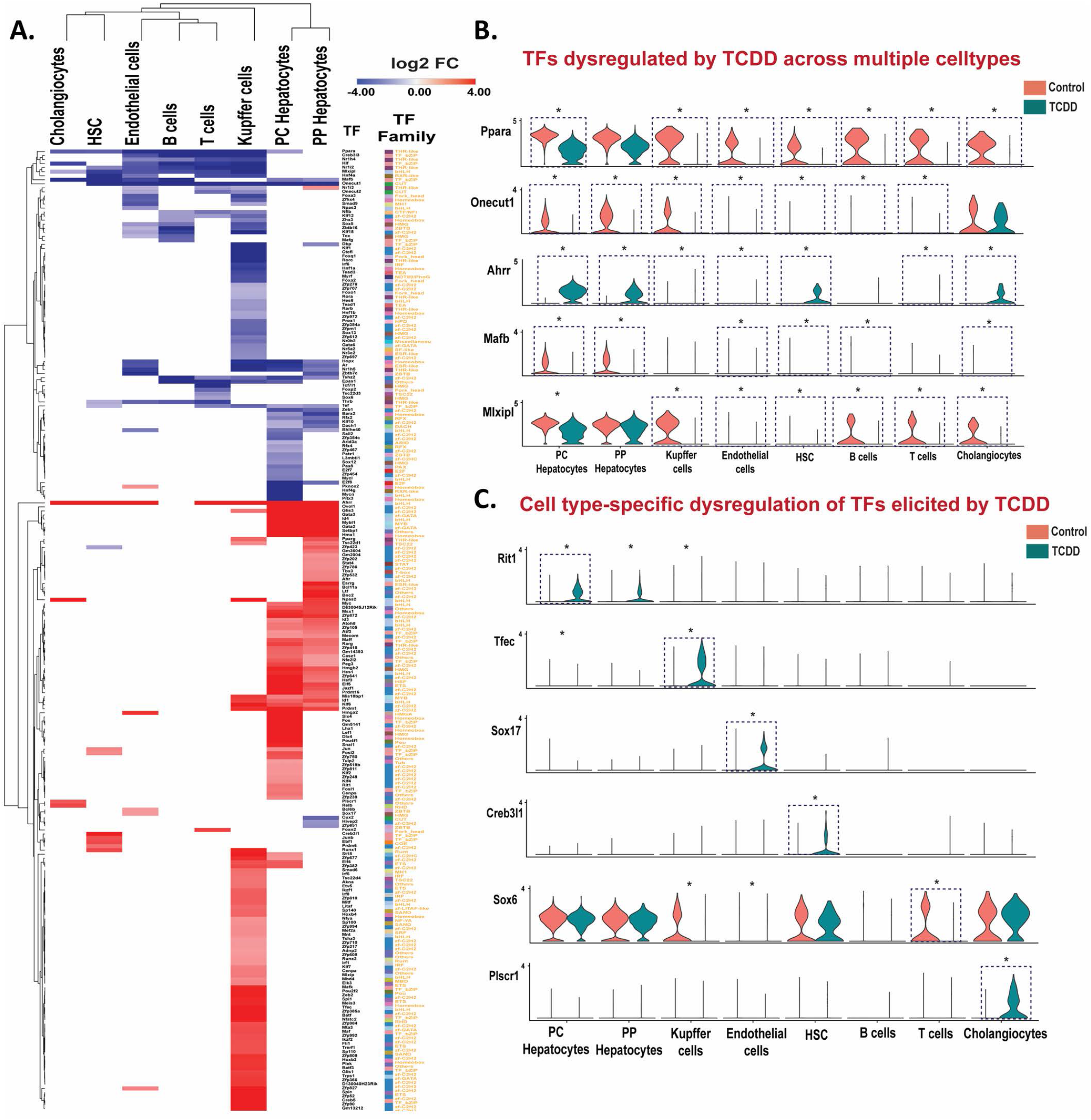
TCDD dysregulates TF expression across multiple liver cell types. **A**. Heat maps of differentially expressed TFs in the indicated cell clusters. The colored cells in the heat map (red or blue) show log2 fold-change (FC) values for TFs whose expression is significantly (FDR <0.05) increased (red) or decreased (purple) by TCDD. **B, C**. Violin plots showing expression data for TFs significantly dysregulated by TCDD (*) in multiple liver cell types (B) or in a cell type-specific manner (C). Blue dotted box with * marks TFs differentially expressed at |fold-change| >4 and FDR <0.05. Also see Table S3C.

TCDD increased expression of the AhR repressor AhRR in all liver cell types, except B cells, with strongest increase seen in hepatocytes (log2 fold-change: 10-11). AhRR binds to the AhR heterodimerization partner Arnt to initiate a negative feedback loop regulating AhR function and TCDD hepatotoxicity ^74^. Five other TFs showed strong up regulation in both hepatocyte populations and in Kupffer cells, including Id1, which promotes HCC proliferation ^75^, and Klf6, a regulator of lipid homeostasis that is up regulated in models of liver injury and activates autophagy in hepatocytes ^76^ (Table S3C). TFs whose expression was dysregulated by TCDD in primarily one liver cell population include Tfec, a macrophage-specific TF ^77^ that was induced 16-fold in Kupffer cells, and Sox17, a regulator of liver lipid metabolism with a role in endothelial regeneration during vasculature injury ^78, 79^, which was specifically induced in endothelial cells by 4-fold. Creb3l1 (Oasis), a key regulatory TF for liver fibrosis-related genes ^80^, was specifically induced, by 15-fold, in hepatic stellate cells, which may contribute to TCDD-induced liver fibrosis. Finally, TCDD induced, specifically in Kupffer cells, Irf1,Irf5 and Irf8, members of the IRF (Interferon regulatory factor) family regulating innate and adaptive immune responses ^81^, while a fourth family member, Irf6, was repressed (Table S3C).

### TCDD-elicited alterations in cell-cell communications

We investigated the effects of TCDD on intrahepatic cell–cell communication patterns, as judged by changes in expression of ligands and their cognate receptors in each liver cell cluster. These analyses were implemented using CellChat and its manually curated list of mouse ligand–receptor interactions ^29^. Strikingly, we observed a large increase in the number of inferred ligand receptor interactions following TCDD exposure (Fig. 5A). These interactions were grouped into a total of 52 signaling pathways, 47 of which have higher overall communication probability (greater relative information flow) in TCDD-exposed liver and five in control liver (Fig. 5B, Table S4). By comparing the overall communication probability between control and TCDD liver, we identified 22 signaling pathways that were specifically active in TCDD-exposed liver (Fig. 5B, Fig. 5C, Table S4). Important signaling pathways altered by TCDD include: increased TGFβ signaling, which promotes the development of NASH in hepatocytes and mediates hepatic stellate cell activation, resulting in a wound-healing response and extracellular matrix deposition ^82, 83^; increased signaling via the adhesion molecule VCAM1 to Kupffer cells from several NPCs, including cholangiocytes, which contributes to the persistence of liver inflammation ^84^ and promotes NASH ^85^; B cell signaling by CD22, a regulator of B cell proliferation ^86^; and signaling from endothelial cells by the pro-inflammatory cytokine IL-1, which contributes to the hepatotoxic and tumor promoting effects of TCDD ^87^.

**Fig. 5.**
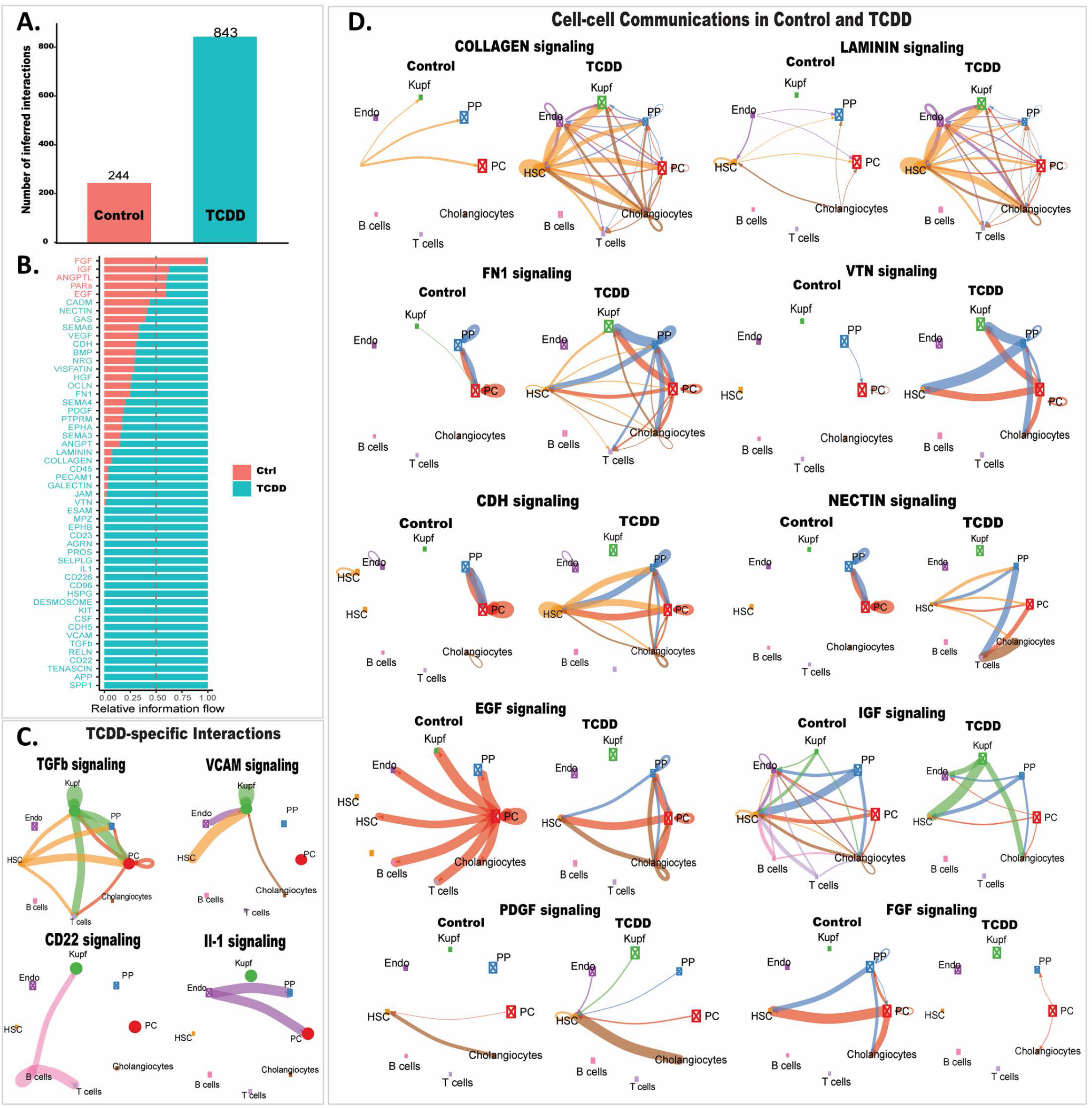
TCDD-induced changes in liver intercellular communication. **A**. Bar plot showing number of ligand-receptor interactions identified by CellChat in snRNA cell clusters from control and TCDD-exposed liver. **B**. Ligand-receptor interactions identified in each dataset are grouped into 52 signaling pathways. X-axis shows the relative information flow of each signaling pathway (Y-axis), identifying intercellular signaling pathways whose activity increases or decreases between control and TCDD-exposed liver. **C**. Circle plots showing examples of TCDD-specific intercellular communication patterns. Edge weights are proportional to the interaction strength: a thicker edge line indicates a stronger signal, while circle sizes are proportional to the number of cells in each cell type. **D**. Circle plots showing examples of differential intercellular communication patterns between control and TCDD-exposed liver. Also see Table S4.

Thirty of the 52 intercellular communication pathways were shared between control and TCDD-exposed liver (Fig. 5B), but often with large differences between the two conditions (Fig. 5D). Thus, major increases in extracellular matrix-receptor interactions were seen in TCDD-exposed liver cells, including signaling via multiple collagens and laminins, and by fibronectin (FN1) and vitronectin (VTN). Increases in intercellular communication involving adhesion proteins, such as nectins and cadherins (CDH), was also found, consistent with AhR regulation of cell adhesion and matrix remodeling ^88^. PDGF, which promotes hepatic stellate cell proliferation and characterizes the fibrotic niche in human liver ^89, 90^, showed increased intercellular connectivity to hepatic stellate cells in TCDD-exposed liver. In contrast, intercellular connections decreased for several growth factor signaling pathways, including EGF, IGF and FGF, which may contribute to the pathological effects of TCDD ^91, 92^.

### Network-essential regulatory lncRNAs in control and TCDD-exposed liver

We implemented gene co-expression network analysis using bigSCale2 ^30^ to construct gene regulatory networks that can be used to identify lncRNAs whose expression is closely associated with specific biological functions. We discovered 5 functional gene modules in both control and TCDD-exposed liver; each module was enriched for distinct sets of pathways and biological functions, as indicated (Fig. 6, Table S5A). Next, for each network, we identified network-essential genes (nodes marked with gene names in Fig. 6) based on four network centrality metrics, namely, Betweenness, PageRank centrality, Closeness, and Degree ^30^, which each serve as a proxy for a gene’s influence on the network (Table S5B). Strikingly, 41 lncRNAs occupied network-essential nodes in the control network and 66 lncRNAs in the TCDD-exposed liver network, of which 6 lncRNAs were identified as essential genes in both networks: Hnf4aos (lnc1966*), Gm36251 (lnc8349), Gm32063 (lnc23914*), AC118710.3 (lnc12772), 0610005C13Rik (lnc6166*), and LOC102632463 (lnc979*) (Fig. S5; Table S5B). Interestingly, 21 of the 41 control network-essential lncRNAs were dysregulated by TCDD, as were 44 of the 66 TCDD network-essential lncRNAs, many of which were down regulated by TCDD in Kupffer cells and/or in endothelial cells (Table S5B, column AI). Five TCDD network-essential lncRNAs (3 with human orthologs) showed robust (>30-fold) induction by TCDD, specifically in periportal and pericentral hepatocytes: lnc12630*, lnc12633, lnc17289, lnc26884* and lnc40479*. The liver regulatory networks were validated by the striking enrichments that the gene targets of the network-essential IncRNAs showed for specific biological pathways. Thus, individual liver network-essential lncRNA regulators were linked to gene ontology terms such as cell adhesion, cell migration and extracellular matrix in the control liver network, and to terms such as fatty acid metabolic process, peroxisome and xenobiotic metabolic process in the TCDD-exposed liver network (Table S5C, Table S5D).

**Fig. 6.**
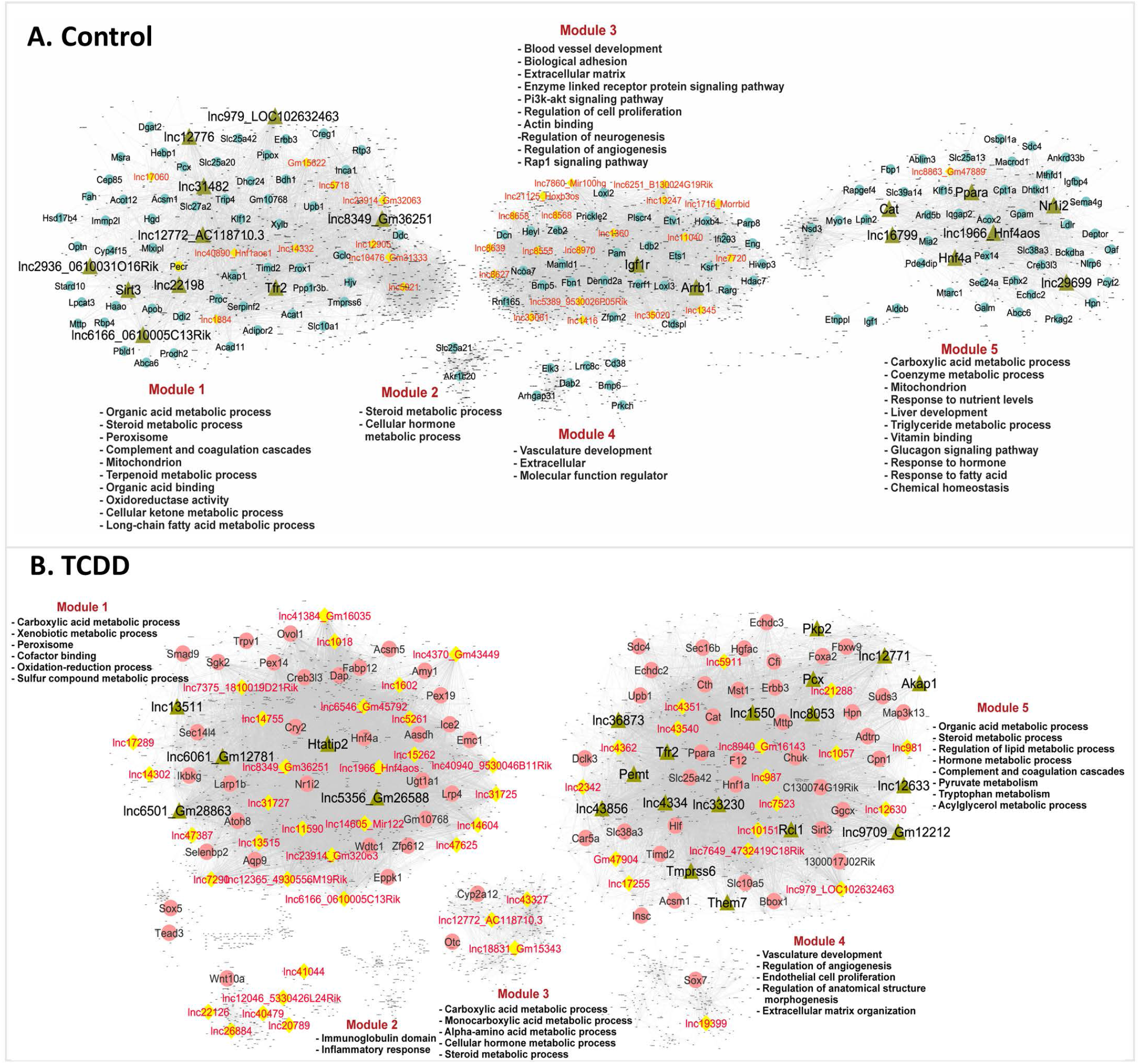
Gene regulatory networks for control and TCDD-exposed mouse liver. Shown are bigSCale2 networks for control **(A)** and TCDD-exposed **(B)** livers, where protein-coding genes or lncRNAs are nodes and the edges between them are correlations based on an adaptive threshold. All nodes identified by gene names represent network-essential genes predicted based on top network metrics. Circular nodes represent protein-coding genes; yellow diamonds indicate lncRNAs; and green triangles identify master regulators, defined as nodes (genes) that have high network centrality metrics calculated in subnetworks extracted from all top 100 ranked protein-coding gene nodes or top 50 ranked lncRNA nodes. Each liver network is subdivided into gene modules that are enriched for the biological functions shown. See Table S5 and https://tinyurl.com/snTCDDliverNetworksKarriWaxman for complete datasets.

Next, we recalculated network metrics using a subnetwork comprised of all top network-essential genes to identify putative master regulators of each network (green triangular nodes in Fig. 6). We identified 19 putative master regulator genes (including 11 lncRNAs) in the control liver network and 22 genes (including 13 lncRNAs) in the TCDD-exposed liver network (Table S5B, Table S5E). Interestingly, one of the lncRNA master regulators of control liver, lnc12772, which is anti-sense to Cyp2d13, showed strong down regulation in all 8 liver cell types in TCDD liver (Table S5B). Importantly, the gene targets of several of the protein-coding gene master regulators showed strong functional enrichment for the known functions of those genes, validating our approach for discovery of network regulatory genes. Examples include the master regulators Nr1i2 (PXR), Sirt3, Ppara and Hnf4a in control liver, and Tmprss6 and Htatip2 in TCDD-exposed liver (Table S5E). Further validation was obtained by IPA Upstream Regulator Analysis, which revealed that 3 of the protein-coding gene master regulators from the control liver network (Nr1i2, Ppara, Hnf4a) are significant Upstream Regulators of their network-determined gene targets (Table S5E).

### Functional clustering of regulatory network-essential lncRNAs

Protein-coding genes that were targets of the network-essential lncRNAs from the control and TCDD-exposed liver networks were input to Metascape ^44^ in a cross-comparison analysis to identify lncRNAs whose network-predicted gene targets share common biological pathways. Network-essential lncRNAs from the control liver network clustered into three major functional groups (Fig. S6), one of which was enriched for metabolic processes and lipid homeostasis, while the other two were variously enriched for terms related to collagen biosynthesis, extracellular matrix, cell-cell adhesion and other non-metabolic processes. Similarly, we identified three clusters of TCDD network-essential lncRNAs, which were distinguished by the functional enrichment terms of their gene targets (Fig. S7). Two of the lncRNA clusters were enriched for both metabolic and non-metabolic processes (angiogenesis, complement cascade), while the third cluster showed strong enrichment for carbon metabolism, xenobiotic metabolism, peroxisome processes and PPAR signaling. Pathways that were either common or specific to the gene targets of the regulatory lncRNAs from each network are shown in Fig. S8.

## Discussion

The use of snRNA-seq for single cell-based transcriptomics has been validated for both mouse and human liver and offers important advantages over single cell-RNA-seq, including the use of flash frozen liver tissue, and the ability to capture nuclei representing all cells present in bulk liver tissue, while minimizing the bias with respect to recovery or viability of individual cell types ^18, 22, 93, 94^ that occurs when using single cells dissociated from liver tissue ^95^. In addition, as a majority of liver-expressed lncRNAs are strongly nuclear-enriched and chromatin bound ^19^, snRNA-seq enables the detection of many more lncRNAs than can be profiled using scRNA-seq analysis ^22^. Here, we show that this approach can be used to characterize thousands of liver-expressed lncRNAs and elucidate their roles in the complex transcriptomic responses to chronic TCDD exposure, a widely studied mouse model of environmental chemical-induced hepatotoxicity and advanced liver disease ^7-9^. We identified more than 4,000 lncRNAs whose expression is dysregulated by TCDD in one or more liver cell types, including 684 lncRNAs specifically deregulated in NPCs. We also discovered 121 lncRNAs, and 689 protein-coding genes, whose zonated expression in hepatocytes across the liver lobule is disrupted by TCDD, the latter finding being consistent with the major zonal toxicity recently described for TCDD ^28^. Gene regulatory networks constructed from the snRNA-seq expression data identified liver network-essential lncRNA regulators specifically linked to functional gene ontology terms, such as cell adhesion, cell migration and extracellular matrix in control (healthy) liver, and fatty acid metabolic process, peroxisome, and xenobiotic metabolic process in TCDD-exposed liver. These findings highlight the power of snRNA-seq to uncover functional roles for many poorly characterized lncRNA genes in biological pathways impacted by TCDD exposure in hepatocytes and liver NPCs.

The snRNA-seq data used for this study ^18^ was obtained from mice chronically exposed to the AhR agonist ligand TCDD over a 4-wk period, which results in hepatic lipid accumulation and promotes progression to steatohepatitis with liver fibrosis ^9^. We observed major changes in lncRNA expression in pericentral hepatocytes, where AhR, the receptor for TCDD, is most highly expressed, and also in periportal hepatocytes, where basal expression of AhR is very low prior to TCDD exposure, but which respond to TCDD with a 5-fold increase in AhR expression. The latter increase undoubtedly contributes to the large number of TCDD-induced gene responses, including lncRNA responses, seen in the periportal hepatocyte subpopulation. Two TFs from the NR superfamily, Nr1i3 (CAR) and Esrrg, showed the same pattern of low basal expression in periportal hepatocytes combined with strong periportal induction following TCDD-exposure, thereby abolishing the pericentral bias in their zonation seen in healthy liver. Nr1i3 (CAR) and Esrrg are both important regulators of liver metabolic homeostasis and responses to liver injury ^55, 96, 97^ and their induction in periportal hepatocytes may be part of the injury response to chronic TCDD exposure. TCDD also induced widespread gene responses in various NPCs, consistent with ^18^, despite the low expression of AhR in those NPCs even after TCDD exposure. Many of the latter gene expression changes are likely to be indirect, downstream effects of TCDD associated with the widespread liver pathology that TCDD induces in this exposure model ^9^.

Many genes active in xenobiotic metabolism and other metabolic processes are regulated by members of the NR superfamily, most notably those NRs that act as xenosensors ^98, 99^. These NRs crosstalk with each other and with TFs that regulate processes such as bile acid synthesis, lipid metabolism and liver fibrosis and inflammation ^100^. We mined the liver snRNA-seq datasets to determine both the basal and the TCDD dysregulated transcriptomic profiles of all mouse NRs across all 8 liver cell types. Expression of 18 of the 49 NRs was undetectable, which in some cases could be due to intrinsic sensitivity limitations of the single cell sequencing technology. In contrast, NRs with established functions linked to lipid metabolic processes and liver disease, most notably Ppara, Nr1h4 (FXRα), Thrb and Rora, were expressed at high levels in both hepatocytes and various NPCs, including hepatic stellate cells, endothelial cells and immune cells. Strikingly, TCDD suppressed the expression of three of these NRs (Ppara, Nr1h4 (FXRα), Thrb) in several NPC populations, including Kupffer cells, which undergo extensive changes following chronic TCDD exposure ^18^ and more generally during NASH pathogenesis ^101^. The strong repression of Ppara in TCDD-exposed liver across all major liver cell types is consistent with an earlier study ^66^ and with the finding that reduced Ppara expression promotes steatosis and NASH development ^102^. These observations, together with our finding that TCDD dysregulates large numbers of other liver TFs (Fig. 4), give important insights into underlying mechanisms for the transcriptional crosstalk between AhR and other TFs, including many NRs, and may help elucidate some of the underlying complexities of TCDD hepatotoxicity.

We investigated the impact of TCDD exposure on cell-cell communication patterns and signaling crosstalk, as deduced from changes in the expression of ligands and their cognate receptors in each liver cell population ^29^. Ligand-receptor interactions within and between liver cell clusters were grouped into signaling pathways, enabling us to capture changes in intra-hepatic cell-cell communication patterns induced by TCDD. Key features of the responses to TCDD include an increase in extracellular matrix and collagen-related signaling pathways and increased PDGF signaling, which are both strong indications of hepatic stellate cell activation leading to liver fibrosis ^89^, a hallmark of chronic exposure to TCDD ^103, 104^. Changes in cell-cell signaling patterns linked to fibrosis and other TCDD-induced hepatotoxic responses include increased TGFβ signaling, which promotes the development of NASH ^82, 83^, increases in signaling by multiple collagens and laminins, which contribute to liver fibrosis, and decreases in intercellular connections for several growth factor signaling pathways associated with pathological effects of TCDD, including EGF, IGF and FGF ^91, 92^. A potential limitation of this approach is that the use of transcriptomics to quantify changes in expression of ligand–receptor pairs does not always correlate with proteomic data ^105, 106^. Increased confidence for such predictions can be obtained by integrating information from proteomics-based technologies that combine scRNA-seq with intracellular protein measurements to profile signaling activity ^107^ or that use mass spectrometry to detect ligand–receptor interactions and assay changes across conditions ^108^.

Single-cell expression data are especially well suited for computing gene regulatory networks, as they do not average out biological signals, unlike bulk transcriptomic data. We used the snRNA-seq datasets to construct gene regulatory networks, which enabled us to associate individual lncRNAs with functional network modules. Further, we used network centrality metrics ^30^ to identify network-essential genes (putative regulatory genes) that drive various biological processes in the liver. We observed only partial overlap between the functional modules identified in the healthy liver (control) and the TCDD-exposed liver networks, consistent with the extensive changes in gene expression and gene regulation that accompany TCDD hepatotoxicity and disease development. The overall network density was higher in the TCDD network (Edges/Nodes=18.1) as compared to the control liver network (Edges/Nodes=8.5) (Table S5G), indicating increased gene regulatory interactions. The network-essential protein-coding genes identified included three major, well-established liver gene transcriptional regulators, Nr1i2, Ppara, Hnf4a, which in an entirely independent approach were verified to be significant Upstream Regulators of their network-determined gene targets by IPA analysis. These and our other findings support the utility of our gene regulatory networks and validate the overall approach described here for discovery of lncRNA genes that have high regulatory potential, and likely high relevance to environmental chemical exposure-induced tissue pathologies and hepatic disease.

## Abbreviations

AhR: aryl hydrocarbon receptor
DEG: differentially expressed gene
GTF: Gene Transfer Format
lncRNA: long non-coding RNA
lnc: followed by a number: numbering system for set of 48,261 mouse liver expressed lncRNAs, where an asterisk (*) marks mouse liver lncRNAs with a human ortholog
NPC: non-parenchymal cell
PC: principal components
scRNA-seq: single cell RNA sequencing
TCDD: 2,3,7,8-tetrachlorodibenzo-*p*-dioxin
TGI: transcript-genome identity
TTI: transcript-transcript identity
UMAP: Uniformed Manifold Approximation and Projection
UMI: unique molecular identifier
VSMC: vascular smooth muscle cell.

## Data availability

Supplementary figures and tables are available online at the journal’s website. A Loupe Browser file, Liver_Control_TCDD_GSE148339, is available at XXX and can be used to query cell clusters in the UMAP shown in Fig. S1 for individual genes, including lncRNA genes, and to perform further downstream data analysis.

## Figure legends

**Fig. S1.**
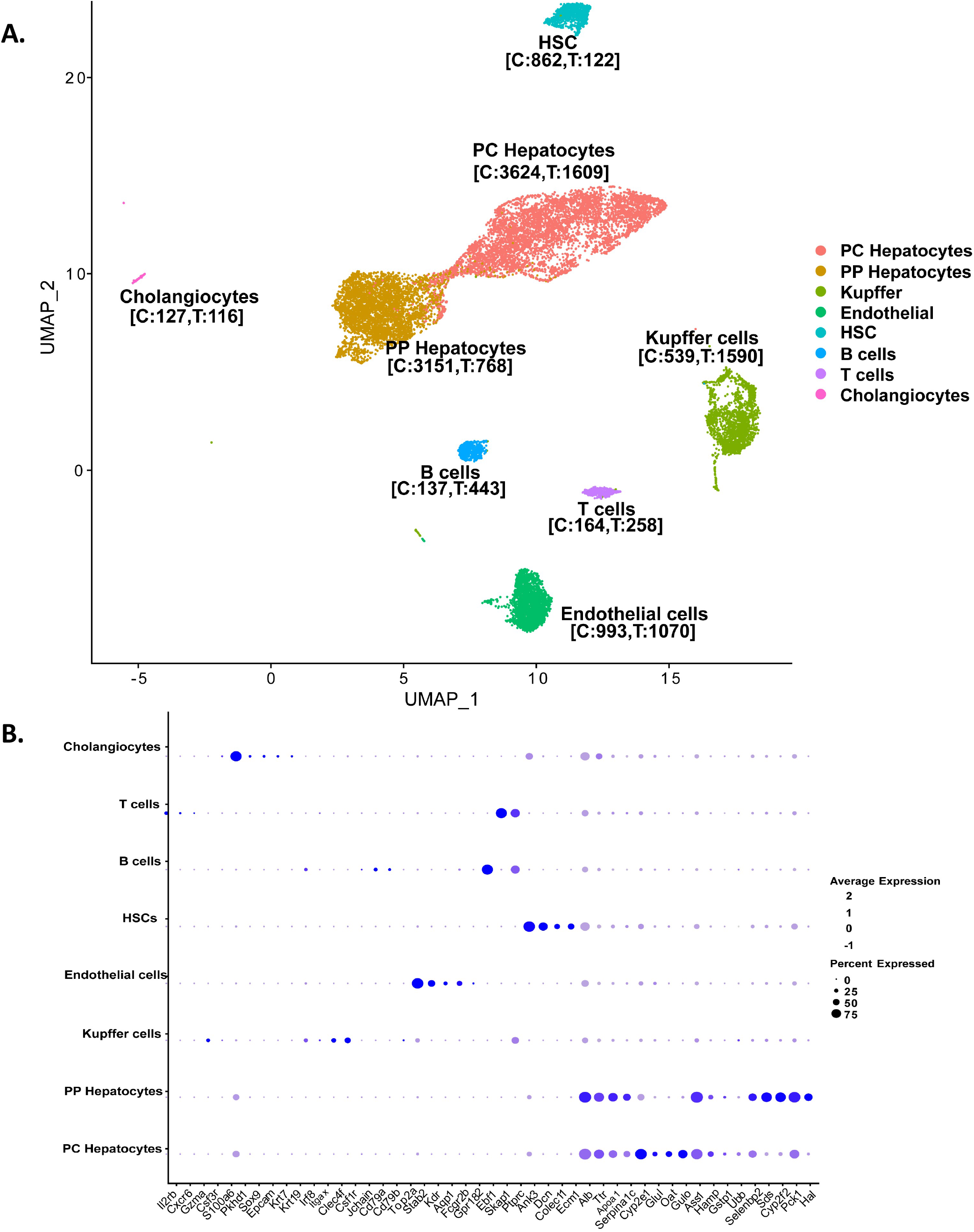
A. Mouse liver cell subpopulations and cell-type specific markers. **A**. UMAP displaying clusters of liver nuclei clusters based on 15,573 single-nuclei transcriptomes integrated from all four snRNA-seq samples, two control liver and two TCDD liver snRNA-seq samples (biological replicates). Cell count values for each cell type and for each condition (C: control; T: TCDD) are indicated in square brackets. **B**. Dot plot showing average expression values for marker genes (shown on X-axis) across the eight hepatic cell clusters (identified on y-axis) for liver nuclei aggregated from all 4 samples -- control and TCDD.

**Fig. S2.**
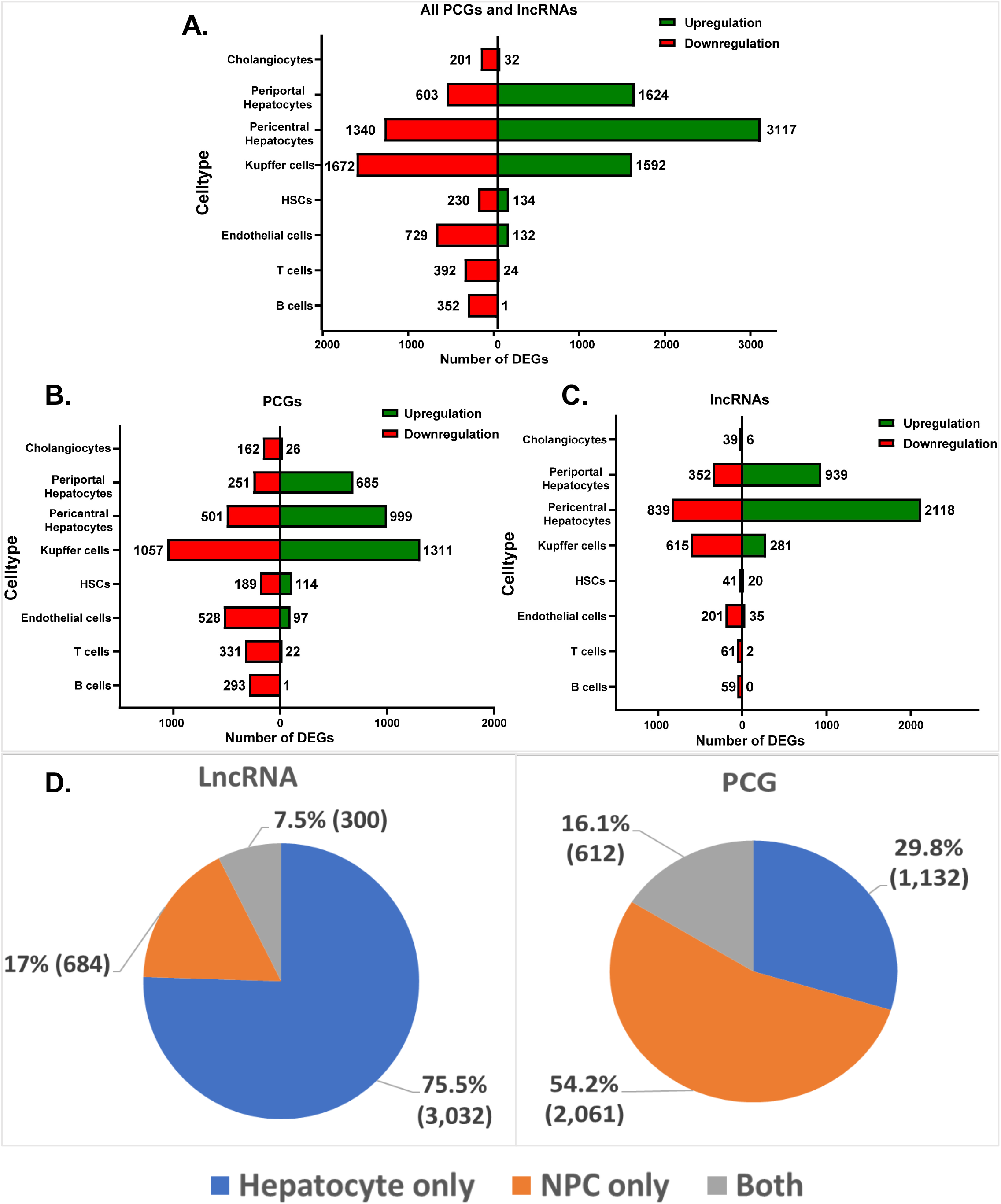
Gene induction and gene repression responses to TCDD. Shown are bar plots indicating the total number of up regulated (green) and down regulated genes (red). (A), all genes, (B) protein-coding genes, and (C) lncRNA genes, in each of the 8 indicated liver cell types, at |fold-change| >4 and FDR <0.05. Many fewer genes were up regulated than were down regulated in several of the NPC clusters. (D) Pie charts showing numbers of lncRNAs and protein-coding genes that are differentially expressed in TCDD-exposed vs control liver in hepatocytes only, in non-parenchymal cells (NPCs) only, or is both hepatocytes and NPCs based on data shown in Table S1, columns AO and AP.

**Fig S3.**
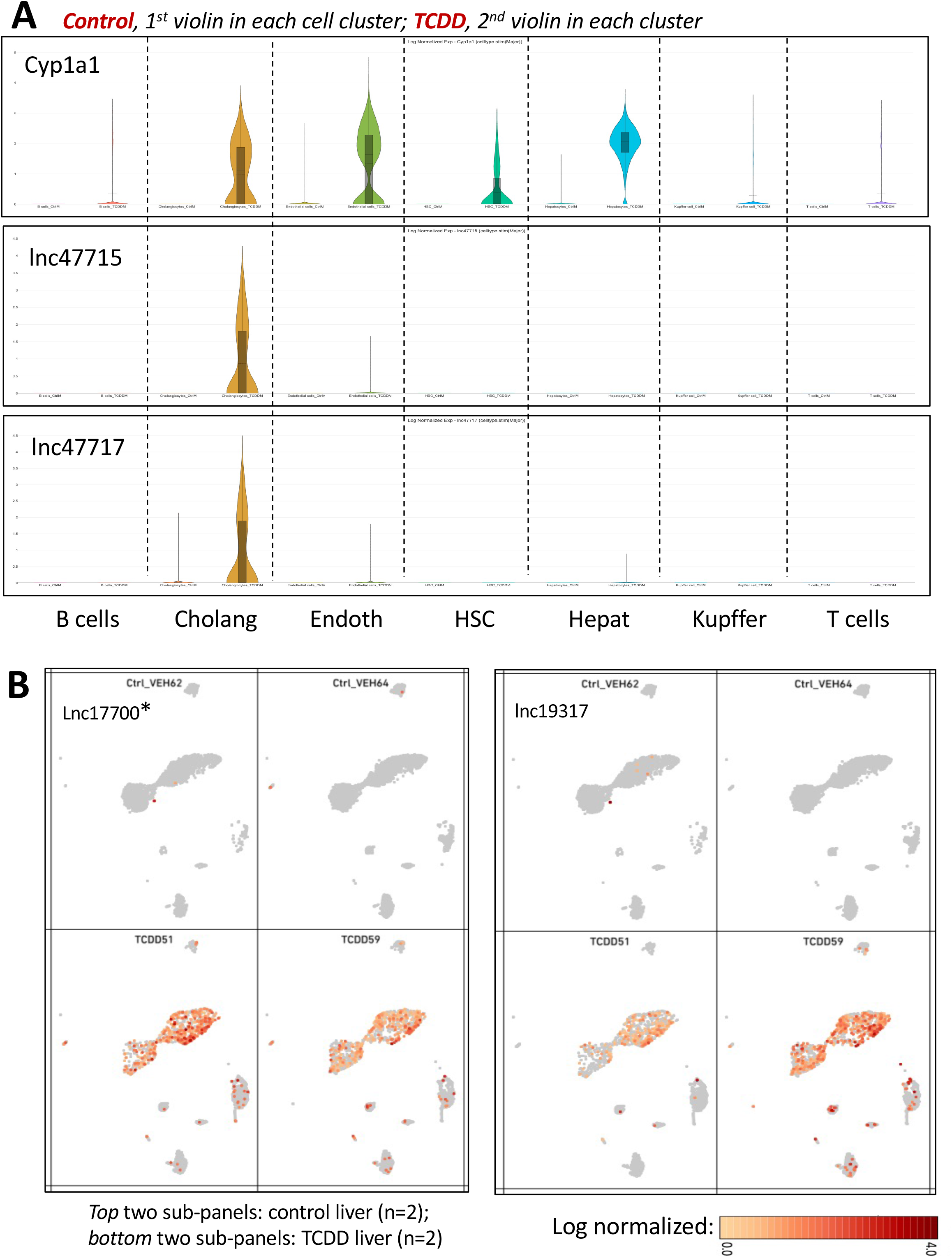
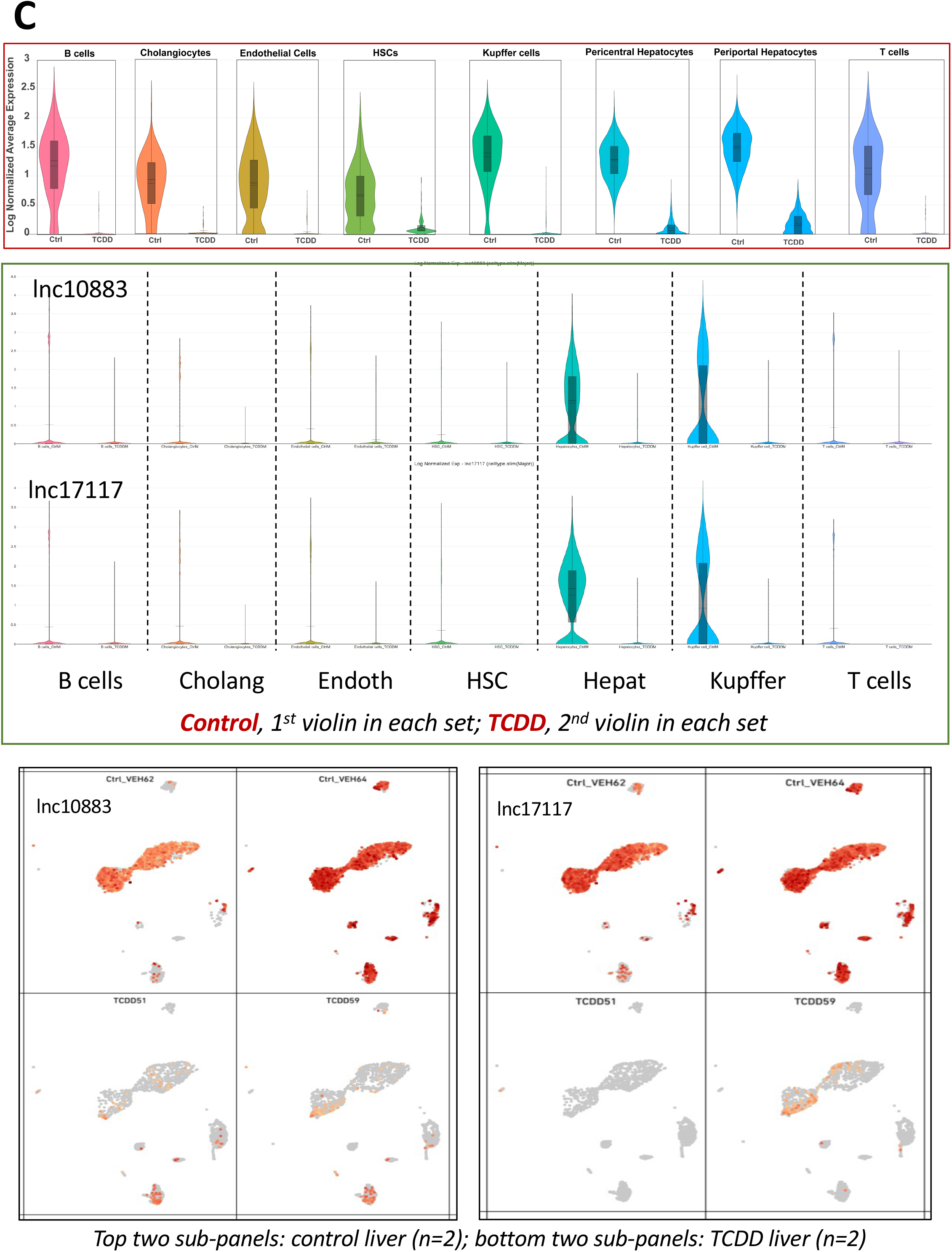
Violin plots and feature plots showing expression data for select TCDD-responsive genes. **A**. Violin plots for Cyp1a1, a classic TCDD/Ahr response gene that is strongly induced by TCDD in multiple liver cell types, and for two lncRNAs whose expression is highly inducible in cholangiocytes but not in other liver cell types. **B**. Feature plots showing expression data for two lncRNAs that are highly induced by TCDD in hepatocytes but not in other cell clusters. Expression data is presented superimposed on the UMAP of cell clusters shown in Fig. S1A, where the individual cell clusters are identified. **C**. Expression data for lncRNAs repressed by TCDD. **Top** panel of C, violin plots showing combined average expression values for 13 lncRNAs repressed by TCDD across multiple liver cell types (Table S1C; lnc35408, lnc10883_Serpina3h, lnc12772_AC118710.3, lnc17117_C730036E19Rik, lnc19880, lnc19881, lnc19883, lnc23635, lnc3297_Gm12909, lnc34452, lnc38016, lnc7809_Gm47465, lnc9497). **Middle** panel of C, violin plots for 2 of the 13 lncRNAs, lnc10883 and lnc17117, which reveals these lncRNAs are most highly expressed in hepatocytes and in Kupffer cells. Repression also occurs in the other cell types, but the level of expression in the other cell types is too low for visualization on the scale of these violin plots. **Bottom** panel of C, feature plots, as described in panel B, for the same two lncRNAs shown in the middle panel of C.

**Figure S4.**
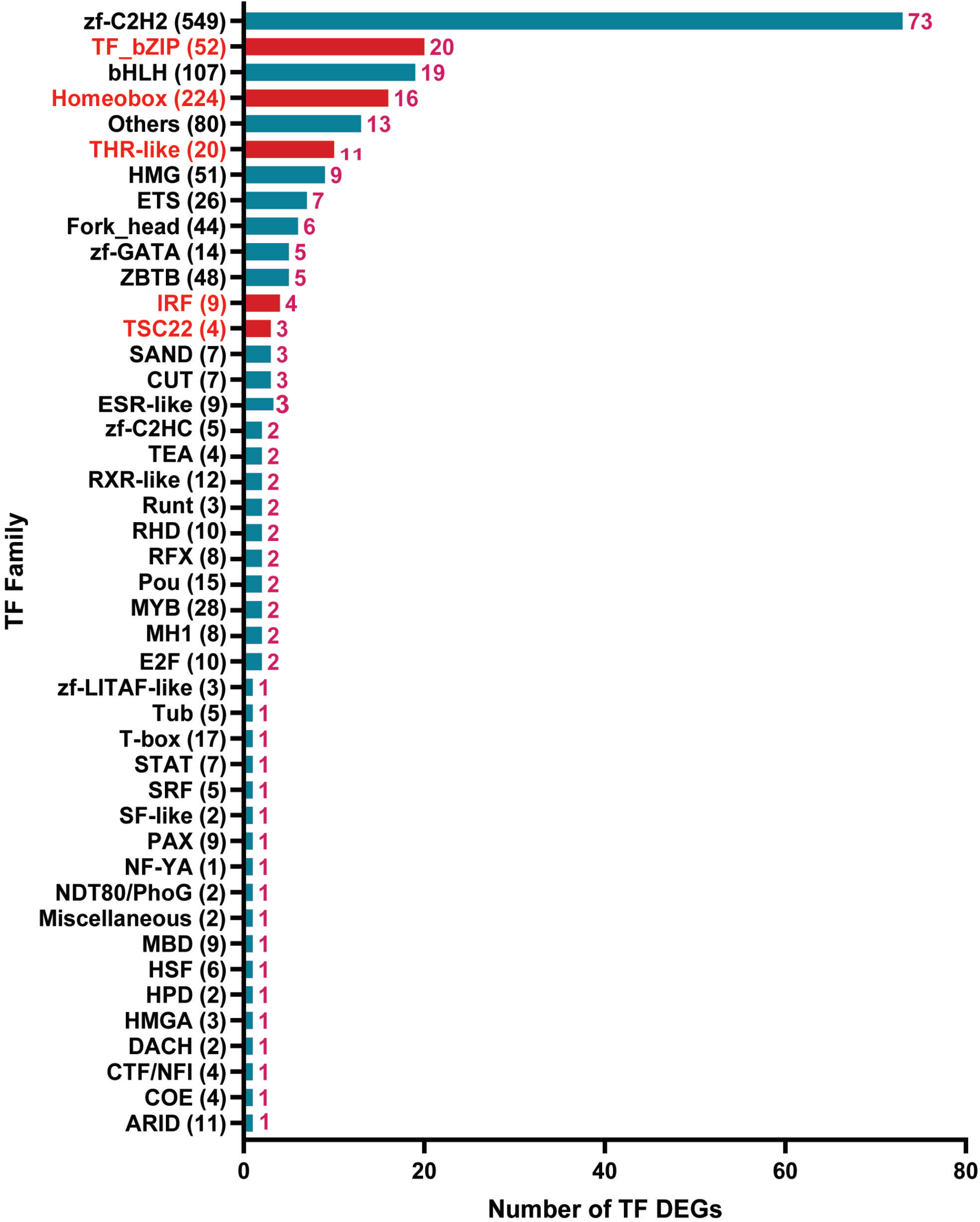
Transcription factor (TF) superfamilies dysregulated by TCDD exposure. Bar plot showing 72 TF families comprising a total of 238 differentially expressed TFs (one TF, Nr4a1, was excluded as it did meet significance in the analysis of separated hepatocyte populations presented in Table S3C; see note in that table legend). TF family classification was based on the DNA-binding domain. *Red* bars mark TF families that were significantly enriched compared to a background set of TFs from the full AnimalTFDB list (Table S3B) that are not differentially expressed at p <0.05, based on Fisher’s exact test. Numbers above each bar, numbers of TFs in the family whose expression was altered by TCDD. The total number of TFs in each family is shown in parentheses along the Y-axis. The THR-like family includes 20 of the 49 NRs shown in Fig. 3.

**Figure S5.**
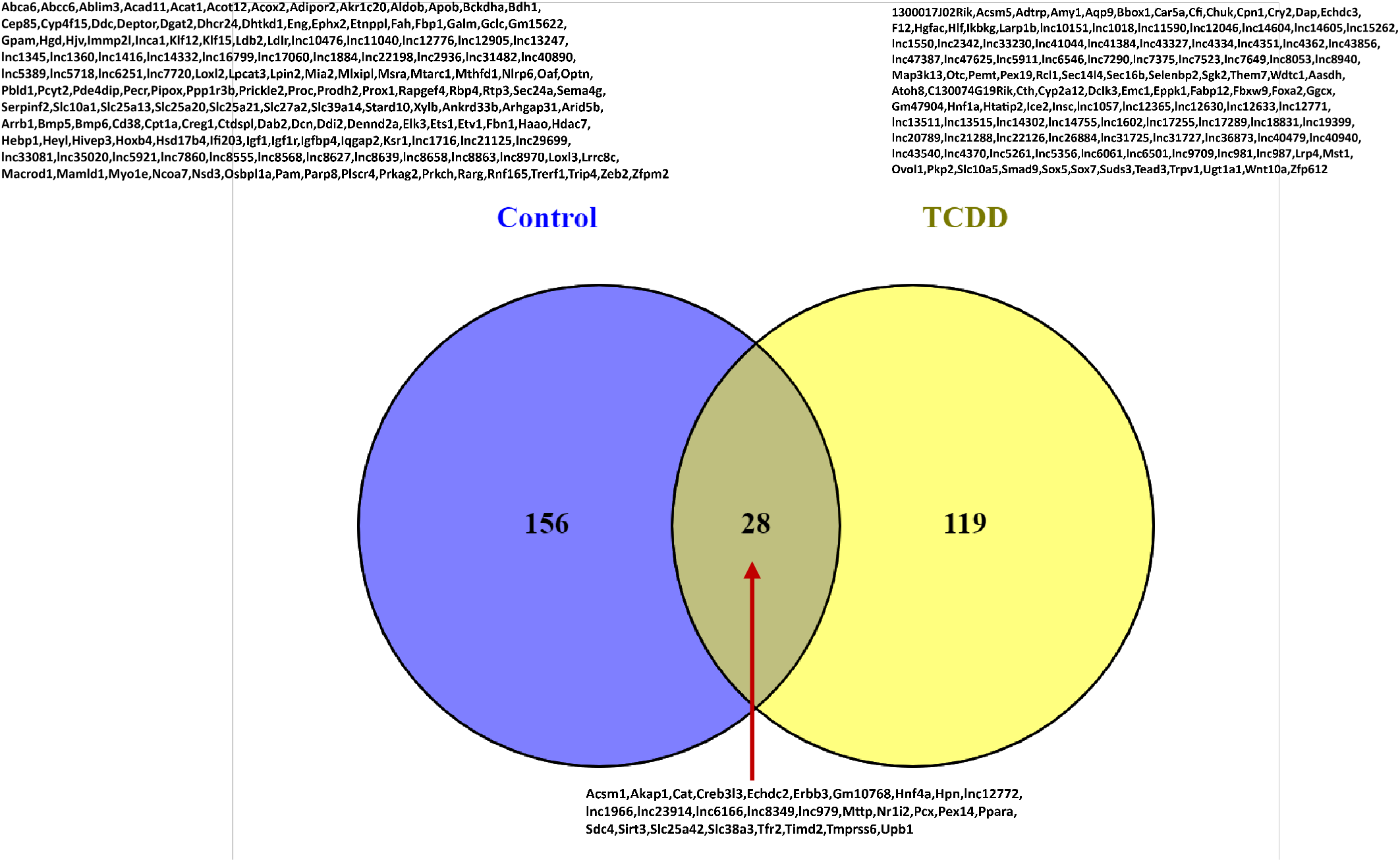
Venn diagram of the number of network-essential genes in control and in TCDD-exposed liver networks. See Table S5B for full listings.

**Figure S6.**
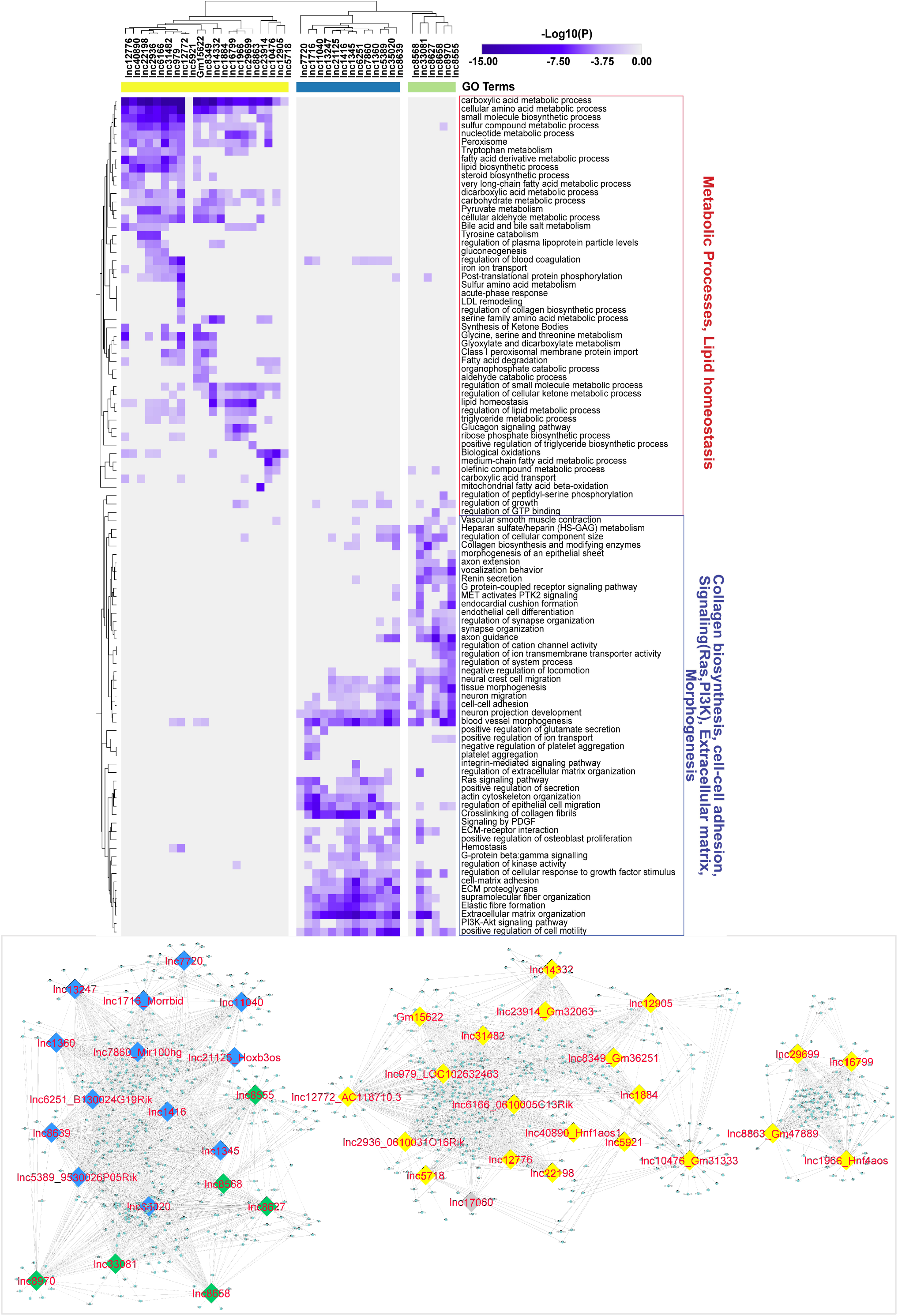
Clustering of 40 network-essential lncRNAs from the control liver gene regulatory network. Protein-coding gene targets that make direct connection with regulatory lncRNAs in the control liver network of Fig. 6A were provided as input to Metascape to obtain top functional enrichment terms based on -log10 (P-value). The color scale represents -log10(P) values, with darker purple coloration indicating more significant p-values. Two-way hierarchical clustering of the functional enrichment heatmap split the 40 lncRNAs into 3 clusters (columns colors) based on their patterns of shared functions. Target genes of lncRNAs from the yellow cluster were enriched for metabolic processes and lipid homeostasis, while the other 2 major functional clusters (blue, green) were primarily enriched in collagen biosynthesis, cell-cell adhesion, signaling and extracellular matrix and morphogenesis, as shown. <u>Bottom</u>: subnetwork of the network shown in Fig. 6A. This subnetwork is comprised of the 40 of the 41 network-essential lncRNAs regulatory lncRNAs and their target genes, which clustered to give gene modules that mirror the clustering results obtained for the same lncRNAs based on their functional group enrichments, shown on top. [Note: the gene targets of 1 of the 41 lncRNA did not yield any enriched pathways; it was therefore excluded from the heatmap]. The subnetwork was derived from the network in Fig. 6A by removing all protein-coding gene regulators and their gene targets, and by retaining regulatory lncRNAs and their direct protein-coding genes connections, which were then used as input for the functional enrichment analysis shown at the top. The regulatory lncRNAs (nodes) at the bottom are color coded to match the lncRNA cluster colors shown in the heat map.

**Figure S7.**
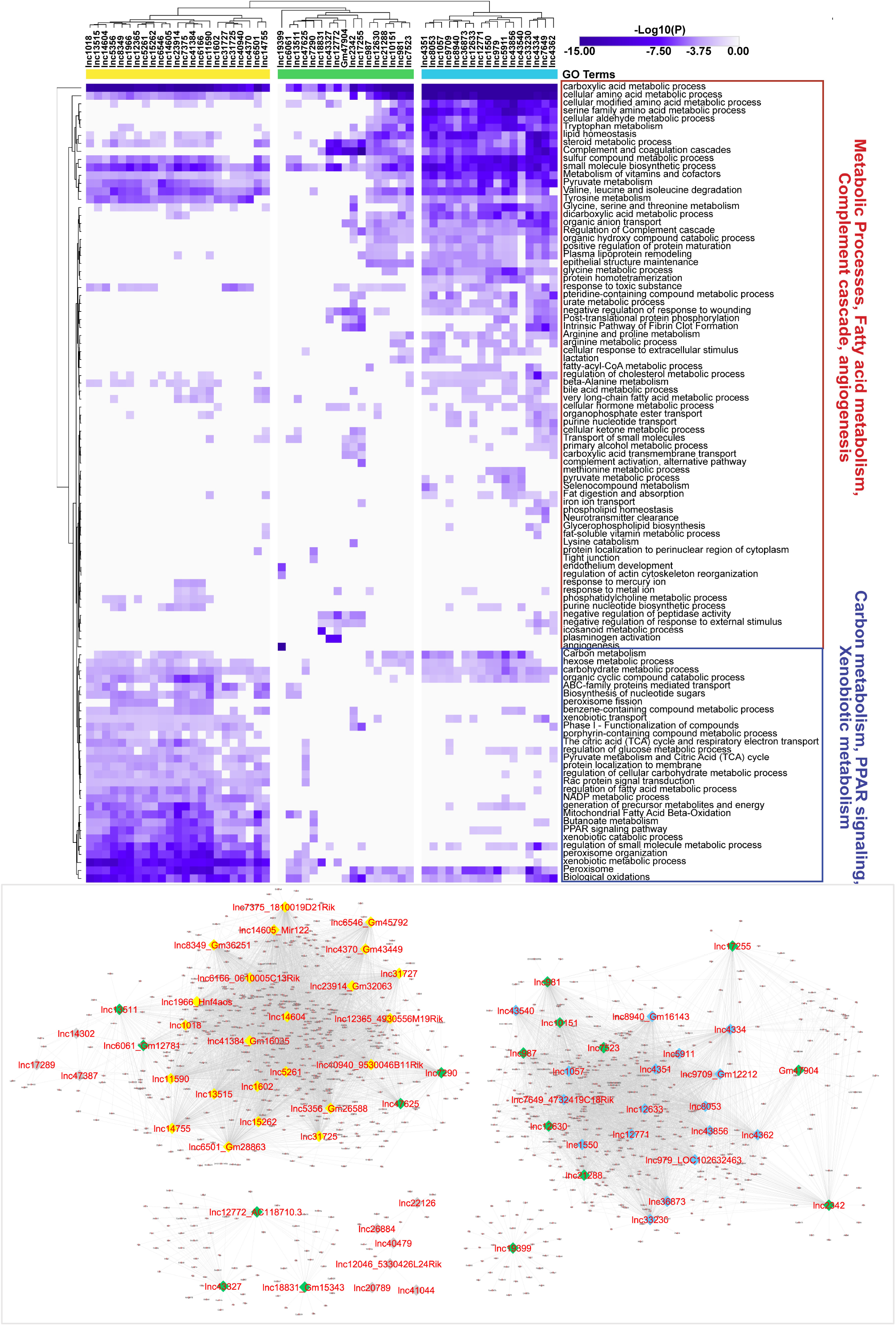
Clustering of 57 network-essential lncRNAs from the TCDD-exposed liver gene regulatory network. Analysis of regulatory lncRNAs from the TCDD network of Fig. 6B, as described in Fig. S6. Two-way hierarchical clustering identified 3 clusters of regulatory lncRNAs, whose target genes were enriched for the pathways and biological processes shown at the right. <u>Bottom</u>: subnetwork of the TCDD network shown in Fig. 6B and obtained as described in Fig. S6.

**Figure S8.**
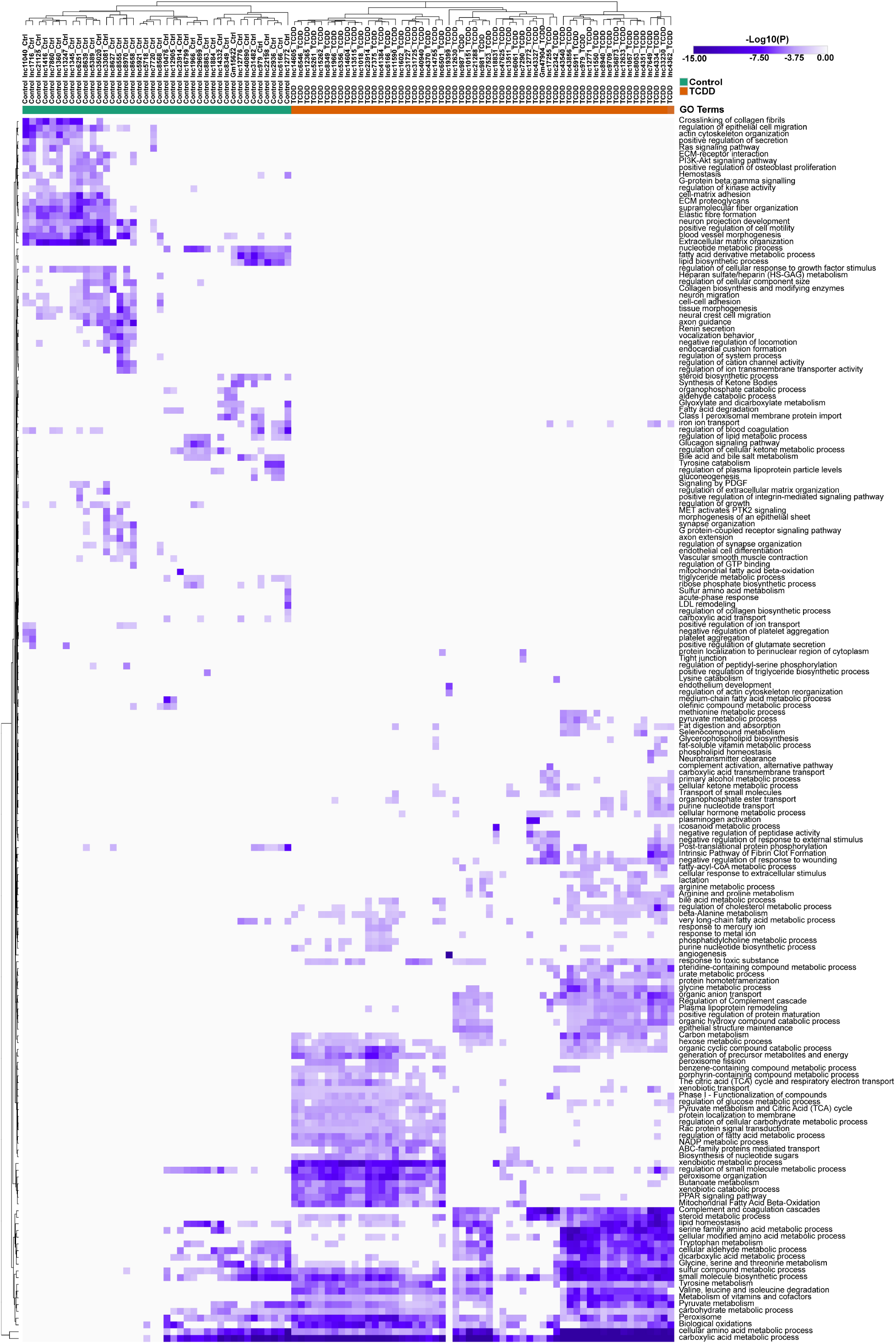
Combined clustering of 97 network-essential lncRNAs from control and TCDD-exposed liver networks. Shown is a merged functional enrichment heatmap showing the top enriched terms (as rows) for genes targets of all regulatory lncRNAs (as columns) from the control and TCDD-exposed liver networks combined and analyzed as described in Fig. S6. This figure indicates that the target genes of the regulatory lncRNAs of the control and TCDD liver networks largely have different enriched functional terms.

